# A protease-mediated mechanism regulates the cytochrome *c*_6_/ plastocyanin switch in *Synechocystis* sp. PCC 6803

**DOI:** 10.1101/2020.07.31.226407

**Authors:** Raquel García-Cañas, Joaquín Giner-Lamia, Francisco J. Florencio, Luis López-Maury

**Affiliations:** Instituto de Bioquímica Vegetal y Fotosíntesis, Universidad de Sevilla- CSIC, c/Américo Vespucio 49, 41092 Sevilla, Spain.; Departamento de Bioquímica Vegetal y Biología Molecular, Facultad de Biología, Universidad de Sevilla, Avenida Reina Mercedes s/n, 41012 Sevilla, Spain; Centro de Biotecnología y Genómica de Plantas UPM – INIA Parque Científico y Tecnológico de la U.P.M. Campus de Montegancedo, Autopista M-40, Km 38, Pozuelo de Alarcón, Madrid 28223, Spain

**Keywords:** Cyanobacteria, plastocyanin, cytochrome *c*_6_

## Abstract

After the Great Oxidation Event (GOE), iron availability was greatly decreased and photosynthetic organisms evolved several alternative proteins and mechanisms. One of these proteins, plastocyanin, is a type I blue-copper protein that can replace cytochrome *c*_6_ as a soluble electron carrier between cytochrome *b*_6_*f* and photosystem I. In most cyanobacteria, expression of these two alternative proteins is regulated by copper availability, but the regulatory system remains unknown. Herein, we provide evidence that the regulatory system is composed of a BlaI/CopY family transcription factor (PetR) and a BlaR membrane protease (PetP). PetR represses *petE* (plastocyanin) expression and activates *petJ* (cytochrome *c*_6_), while PetP controls PetR levels *in vivo*. Using whole-cell extracts, we demonstrated that PetR degradation requires both PetP and copper. Transcriptomic analysis revealed that the PetRP system regulates only four genes (*petE*, *petJ*, *slr0601*, and *slr0602*), highlighting its specificity. Furthermore, the presence of *petE* and *petRP* in early branching cyanobacteria indicates that acquisition of these genes could represent an early adaptation to decreased iron bioavailability following the GOE.

**Significance Statement:** After the appearance of oxygenic photosynthesis, Fe became oxidized and its solubility and availability were greatly decreased. This generated a problem for most organisms since they are strongly dependent on Fe, especially photosynthetic organisms. In response, organisms evolved alternatives to Fe-containing proteins such as plastocyanin, a copper protein that substitutes for cytochrome *c*_6_ in photosynthesis. Expression of these two proteins in cyanobacteria is regulated by Cu availability, but the regulatory system remains unknown. Herein, we describe the regulatory system for these alternative proteins in photosynthesis in cyanobacteria. The mechanism involves a transcription factor (PetR) and a membrane protease (PetP) that degrades PetR in the presence of Cu.

## Introduction

Cyanobacteria are prokaryotes that perform oxygenic photosynthesis, and their appearance on earth led to a dramatic increase in oxygen in the atmosphere. Although the timing of the Great Oxygenation Event (GOE) is well established, its extent is still a matter of dispute (1–4). The GOE caused a massive extinction because oxygen was toxic to most life forms (5, 6) and provoked Fe oxidation (Fe^2+^ to Fe^3+^), which made Fe much less available since Fe^3+^ precipitates. This created an additional problem for all forms of life because Fe was already established as a major cofactor in biochemical reactions (7, 8). In fact, photosynthesis has high Fe requirements and photosynthetic organisms usually have a higher Fe quota than non-photosynthetic organisms because Fe is contained in proteins of both photosystems, in the cytochrome *b*_6_*f* complex, in the soluble thylakoid lumen electron carrier cytochrome *c*_6_ (Cyt*c*_6_), and in ferredoxin outside the thylakoids (9). Nowadays, Fe is a limiting nutrient in most ecosystems, especially in aquatic environments (10). Consequently, photosynthetic organisms evolved several alternatives to iron-containing proteins. In most cyanobacteria and some algae, Cyt*c*_6_ can be replaced by plastocyanin (PC), a type I blue-copper protein that acts as a thylakoid lumen soluble electron transporter. These two electron carriers are present in millimolar concentrations inside the thylakoid lumen (11), where Cyt*c*_6_ is replaced by PC, and this substantially reduces the requirement for Fe. In addition, PC uses Cu which is much more soluble (and therefore available) than Fe in an oxidizing atmosphere. Interestingly, plants have only retained PC for photosynthetic electron transfer (12).

Expression of these two alternative electron carriers is regulated by copper availability in both cyanobacteria and algae (13–15). The regulatory system has been well characterized in the green alga *Chlamydomonas reinhardtii*. CRR1, an SQUAMOSA promoter binding protein-like transcription factor (SBP), activates *CYC6* (encoding Cyt*c*_6_) expression in response to copper limitation (16). CRR1 is able to bind both Cu^+^ and Cu^2+^, with the latter inhibiting the DNA-binding ability of the SBP domain (16). However, the regulatory system has not been identified in cyanobacteria, even though the Cyt*c*_6_/PC switch was discovered more than 40 years ago (13, 14, 18–21), and it is frequently used to control the expression of genes in several model systems (22–24). The switch responds to the presence of copper at the transcriptional level; *petJ* (encoding Cyt*c*_6_) is repressed while *petE* (encoding PC) is induced (18, 21, 25–27). Several regulatory systems that respond to copper have been described in cyanobacteria including CopRS (25, 27), InrS (27, 28), and BmtR (29), but none are involved in regulating these two genes.

Herein, we unveiled the regulatory mechanism for the Cyt*c*_6_/PC switch in cyanobacteria. It involves two genes: one encoding a transcription factor of the BlaI superfamily (PetR) that binds to both *petE* and *petJ* promoters, repressing *petE* and activating *petJ* in the absence of copper. The second gene encodes a protease (PetP) that degrades PetR in response to copper. PetR degradation is activated by copper and requires PetP and copper *in vivo* and in an *in vitro* assay system, suggesting that PetP is responsible for PetR degradation. The system is widespread in all cyanobacteria, including early-branching cyanobacteria such as *Gloeobacter* and *Pseudoanabaena*, with the exception of most strains of the *Synechococcus*/*Prochlorococcus* groups.

## Results

### A BlaI/CopY homolog regulates the *petJ*/*petE* switch in *Synechocystis*

Bioinformatics analysis of genes within the vicinity of *petE* and *petJ* in 169 cyanobacterial genomes revealed that a transcription factor of the BlaI/CopY family (renamed PetR) was associated with *petJ* and/or *petE* genes in 13 genomes (SI Appendix, Figure S1). Although the overall number of genomes with this gene association was low, co-occurrence of *petR* with both *petE* and *petJ* was higher. Specifically, 75% of genomes harboring *petE* and *petJ* also contained *petR* (and a linked gene encoding a BlaR homolog; see below; SI Appendix, Figure S1), and these two genes were only present in genomes containing *petE* and *petJ* (SI Appendix, Figure S1; Supplementary dataset S1) with a few exceptions. This raised the possibility that *petR* could be related to the *petE*/*petJ* switch. In order to test this hypothesis, a mutant in the *slr0240* gene encoding PetR in *Synechocystis* sp. PCC 6803 (*Synechocystis* hereafter) was generated (PETR strain). The mutant strain fully segregated (SI Appendix, Figure S2) and the phenotype was indistinguishable from that of the wild-type (WT) strain on BG11C plates (the standard defined medium for cyanobacteria; see below). Expression of *petJ* and *petE* genes was analyzed by RNA blot in both WT and PETR strains after copper addition to cells grown in BG11C lacking Cu (BG11C-Cu). In WT cells, *petE* was induced in response to copper and *petJ* was repressed (Figure 1A), as previously described (18, 25–27). By contrast, in the PETR strain *petE* was expressed even in the absence of copper and *petJ* was not expressed under any conditions (Figure 1A). Conversely, *copM* expression, which is induced in response to copper (25, 30), was similar in both strains. At the protein level, PC was induced in WT cells upon copper addition, while Cyt*c*_6_ abundance decreased after 4 and 24 h. CopM levels increased after copper treatment, with the unprocessed form of the protein accumulating first, followed by the processed at later timepoints (Figure 1B). By contrast, Cyt*c*_6_ was not detected under any conditions in the PETR strain, and PC was detected even in BG11C-Cu medium (Figure 1B). An increase in PC levels was still evident in the PETR strain after copper addition (Figure 1B), probably due to an increase in copper-bound PC, which is more stable than apoPC (25), since mRNA levels did not change after copper addition (Figure 1A). No changes in the CopM induction pattern were observed (Figure 1B). When an additional copy of *petR* was introduced in the PETR strain, either under its own promoter (PETR2 strain) or under the strong *cpcB* promoter (PETR3 strain), a WT phenotype was restored (SI Appendix, Figure S3). This indicates that *petR* was the only gene responsible for regulating these two genes. Similar results were obtained when protein levels were analyzed in steady-state cells of the WT strain grown in media with differences in copper availability. PC was high in copper-containing medium (BG11C 330 nM Cu), lower in BG11C-Cu (~30 nM Cu), and much lower in BG11C-Cu+ bathocuproinedisulfonic acid (BCSA), a copper-specific chelator (no copper; Figure 1C). The PETR strain expressed PC in all conditions, but accumulated higher levels in copper-containing media. Meanwhile, Cyt*c*_6_ was not detected in the PETR strain in any media, and WT cells displayed higher levels in BG11C-Cu+BCSA than in BG11-Cu (Cyt*c*_6_ was almost undetectable in copper-containing medium; Figure 1D).

**Figure 1.**
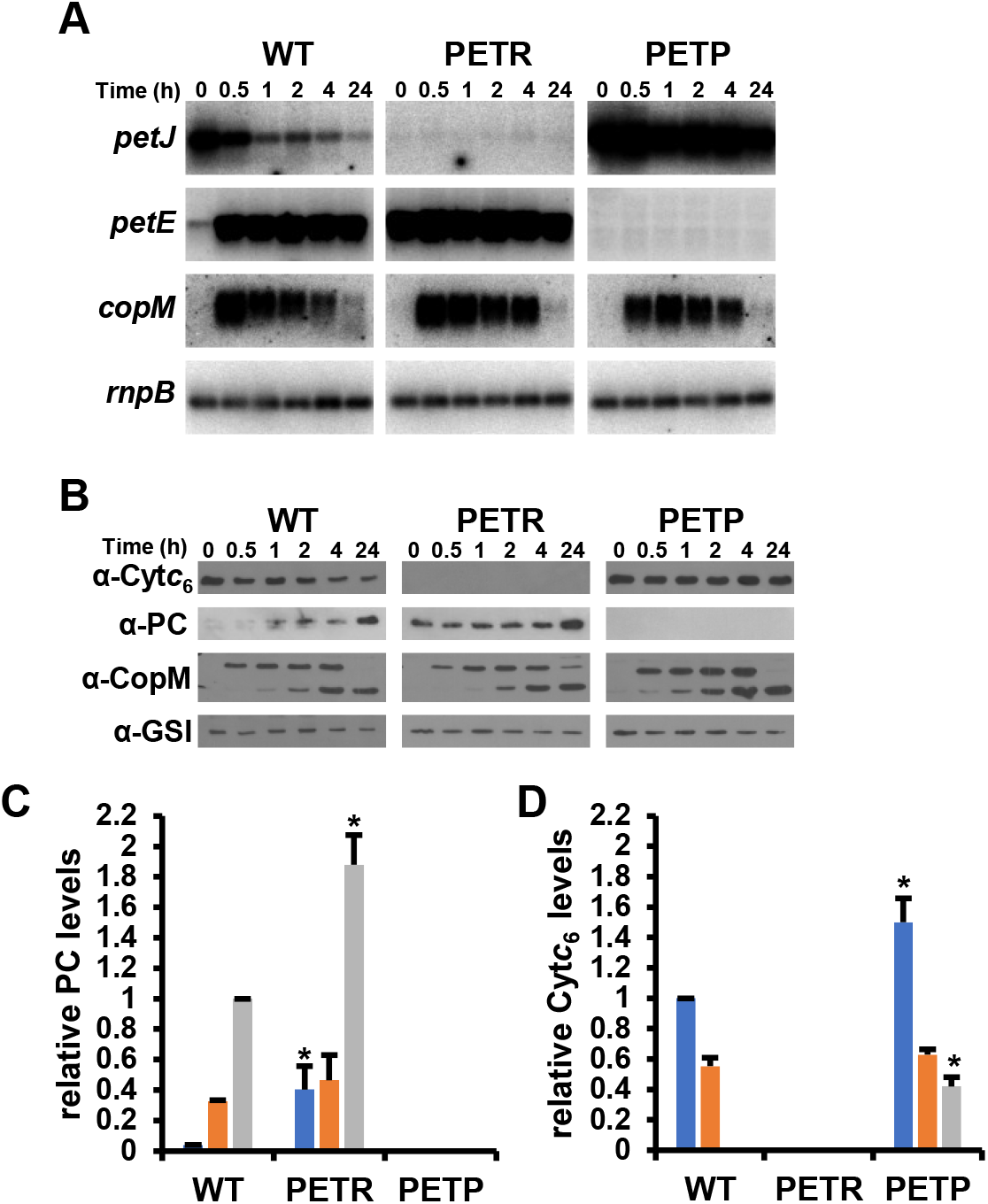
PetRP regulatory system controls *petJ*/*petE* switch in *Synechocystis* sp. PCC 6803. A. RNA blot analysis of *petJ, petE* and *copM* in WT, PETR and PETP strains in response to 0.5 μM copper addition. Total RNA was isolated from cells grown in BG11C-Cu medium at the indicated times after addition of 0.5 μM of copper. The filters were hybridized with *petJ*, *petE* and *copM* probes and subsequently stripped and re-hybridized with a *rnpB* probe as a control. B. Immunoblot analysis of Cyt*c*_6_, PC, CopM and GSI in WT, PETR and PETP strains in response to 0.5 μM copper addition. Cells were grown in BG11C-Cu medium and cells were harvested at the indicated times addition of 0.5 μM of copper. 5 μg of total protein from soluble extracts was separated by 15 % SDS-PAGE and subjected to inmunoblot to detect PC, Cyt*c*_6_,CopM or GSI. C. Quantification of PC levels in WT, PETR and PETP strains grown in BG11C-Cu +BCSA (blue bars), BG11C-Cu (orange bars) or BG11C (grey bars). 10 μg of total soluble proteins were loaded and compared to serial dilutions (100%, 50%, 25% and 12.5%) of the WT extract prepared from cells grown in BG11C. Data are the mean ± SE of three biologically independent experiments. Asterisk indicate significant difference to WT in the same condition (*t*-test; *p*<0.05). D. Quantification of Cyt*c*_6_ levels in WT, PETR and PETP strains grown in BG11C-Cu +BCSA (blue bars), BG11C-Cu (orange bars) or BG11C (grey bars). 10 μg of total soluble proteins were loaded and compared to serial dilutions (100%, 50%, 25% and 12.5%) of the WT extract prepared from cells grown in BG11C-Cu+BCSA. Data are the mean ± SE of three biologically independent experiments. Asterisk indicate significant difference to WT in the same condition (*t*-test; *p*<0.05).

### Slr0241(PetP) is also involved in *petJ*/*petE* regulation

Downstream of *petR* there is another gene (*slr0241*, herein named *petP*) that encodes an M48/M56 superfamily of proteases (Pfam families PF01435 and PF05569). The *petP* start codon overlaps with the stop codon of *petR*, and both genes were expressed as a single transcriptional unit (SI Appendix, Figure S4). Analysis of the PetP sequence using TMHMM 2.0 software (31) predicted four transmembrane helices (residues 4–21, 28–50, 65–87, and 256–278), a 14 residue loop connecting TM2 and TM3, and a cytoplasmic domain (88-256) that contains the HEXXH conserved catalytic motif (SI Appendix, Figure S5). This domain organization is very similar to that of BlaR from *Staphylococcus aureus* (32), which is involved in regulating β-lactam resistance by degrading BlaI, although it lacks the C-terminal periplasmic domain that binds to β-lactams. Furthermore, the organization of *petRP* is conserved in all cyanobacterial genomes (SI Appendix, Figure S1, and Supplementary dataset S1), suggesting that both genes might be functionally related. To test this hypothesis, we mutated the *petP* gene. The resulting mutant strain (PETP) was viable (SI Appendix, Figure S2) and exhibited a similar growth phenotype to the WT strain on BG11C plates (see below). When gene expression was tested in this mutant, *petE* was repressed both in the presence and absence of copper, while *petJ* was expressed constitutively (Figure 1A). The same results were observed at the protein level, with Cyt*c*_6_ detected both in the presence and absence of copper, while PC was undetectable under any conditions (Figure 1B). When steady-state proteins levels were analyzed in different copper-containing media, PC was not detected in the PETP strain in any media, while Cyt*c*_6_ was detected in all media. Despite this, Cyt*c*_6_ levels were lower in copper-containing medium than in media lacking copper (Figure 1D). These results indicate that *petP* is involved in the regulation of both genes, acting in an opposite way to *petR*. A double mutant lacking both *petR* and *petP* genes was also generated (PETRP; SI Appendix, Figure S2). The PETRP strain displayed the same phenotype as the PETR strain (SI Appendix, Figure S6), revealing that *petP* acts upstream of *petR*.

### PetP regulates PetR levels in response to copper

Given that PetP belongs to the M48/M56 superfamily of proteases, this protein might regulate PetR through proteolysis. To investigate this hypothesis, we generated an antiserum against PetR that allowed us to monitor PetR protein abundance in response to copper. In the WT strain, PetR was detected in cells cultivated in BG11C-Cu, and it was rapidly degraded after copper addition, since *petRP* mRNA levels did not change following copper addition (SI Appendix, Figure S4). Two bands were detected (~15 kDa and ~12 kDa, Figure 2A), suggesting an initial and specific cut, and subsequent degradation of the fragments. Indeed, in order to consistently detect PetR, whole cells were loaded onto gels (see Materials and Methods) because PetR was quickly degraded during protein extract preparation. The best studied homologs of PetRP are BlaIR proteins involved in antibiotic resistance from *S. aureus* (32). Processing of these proteins was initially identified in BlaI, it takes places after Phe100 in the amino acid sequence, and is completely conserved in all cyanobacterial PetR proteins (SI Appendix, Figure S7). As expected, the theoretical molecular mass of the fragment generated by this digestion was consistent with the expected size detected by immunoblotting (Figure 2A and SI Appendix, Figure S7; expected sizes of full-length and processed proteins are 15.80 and 12.38 kDa, respectively). In contrast to the WT strain, PetR levels did not change in response to copper in the PETP strain (Figure 2). Therefore, the initial cleavage is probably mediated by PetP, and the fragments are probably further degraded by other unspecific proteases. As expected, PetR was not detected in the PETR strain (Figure 2 and SI Appendix, Figure S3E). When PetR protein abundance was analyzed in steady-state cultures grown in different copper regimes, levels were clearly regulated in the WT strain, since they were higher in the presence of BCSA than in BG11-Cu, and not detected in copper-containing media (Figure 2B and C). PetR levels were higher in the PETP strain than in the WT strain, even when WT cells were cultured in the presence of BCSA, although levels were decreased slightly at higher copper availability (Figure 2B and C). These results are consistent with the Cyt*c*_6_ and PC levels measured under the same conditions (Figure 1C and D), and demonstrate that PetR protein abundance is regulated by PetP *in vivo* according to Cu availability.

**Figure 2.**
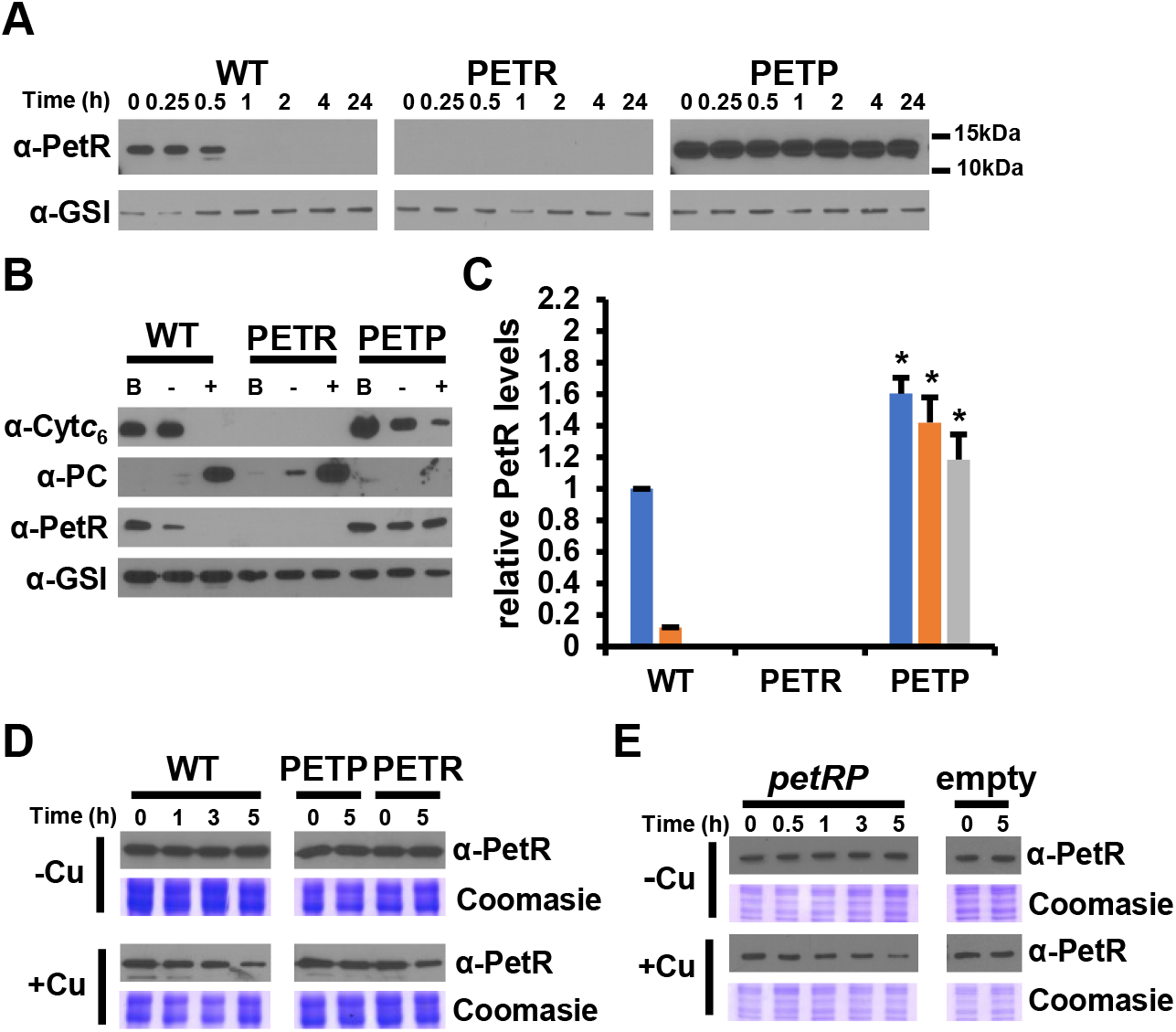
PetP is a copper activated protease that regulates PetR levels. A. Immunoblot analysis of PetR in WT, PETR and PETP strains in response to 0.5 μM copper addition. Cells were grown in BG11C-Cu medium and harvested at the indicated times after addition of 0.5 μM of copper. Whole cells were loaded (0.2 OD_750nm_), separated by 15 % SDS-PAGE and subjected to immunoblot to detect PetR or GSI as loading control. B. Immunoblot analysis of Cyt*c*_6_, PC and PetR in WT, PETR and PETP strains grown in BG11C-Cu+BCSA (B), BG11C-Cu (-) or BG11C. 10 μg of total soluble proteins were separated by 15 % SDS-PAGE and subjected to immunoblot blot to detect PC, Cyt*c*_6_, or GSI as loading control. For PetR whole cell extracts were used and 20 μl (equivalent to 0.2 OD_750nm_) were loaded per lane. C. Quantification of PetR levels in WT, PETR and PETP strains grown in BG11C-Cu +BCSA (blue bars), BG11C-Cu (orange bars) or BG11C (grey bars). 20 μl of the whole cells extract were loaded and compared to serial dilutions (100%, 50%, 25% and 12.5%) of the WT extract prepared from cells grown in BG11C-Cu+BCSA. Data are the mean ± SE of three biologically independent experiments. Asterisk indicate significant difference to WT in the same condition (*t*-test; *p*>0.05). D. GST-PetR was incubated for the indicated time with total extracts prepared from WT, PETR and PETP strains grown in BG11C-Cu. 0.5 μM CuSO_4_ was added to the extracts as indicated. Samples were taken at the indicated times, mixed with Laemmli buffer, boiled, separated in 12% SDS-PAGE gels and GST-PetR was detected using PetR antibodies. E. GST-PetR was incubated for the indicated time with whole cell extracts prepared from *E. coli* carrying pN_petRP or pN_Nat (an empty plasmid) grown in M9 minimal media. 0.5 μM CuSO_4_ was added to the extracts as indicated. Samples were taken at the indicated times, mixed with Laemmli buffer, boiled, separated in 12% SDS-PAGE gels and GST-PetR was detected using PetR antibodies

### PetP degrades PetR directly in a copper-dependent manner

To further characterize this regulation, we performed an *in vitro* degradation assay in which recombinant GST-PetR was incubated with total cell extracts from WT, PETR, and PETP strains, and GST-PetR processing was followed by immunoblotting. GST-PetR was not degraded by whole-cells extracts prepared from WT, PETP, or PETR strains when copper was not added to the extracts (Figure 2D). In contrast, when copper was added to extracts, GST-PetR was degraded by WT and PETR extracts, but not by PETP extracts, indicating that a PetP-mediated PetR degradation was activated by copper (Figure 2D).

To confirm that PetP was directly involved in PetR degradation, and that this degradation was not mediated by an additional protein from *Synechocystis*, *petRP* genes under their own promoter were cloned in a plasmid and transformed into *E. coli*. Total extracts were prepared from cells carrying a plasmid containing *petRP* or an empty plasmid, and degradation of GST-PetR was monitored. The results were similar to those obtained using total extracts from *Synechocystis*: GST-PetR was degraded in total extracts prepared from cells containing a plasmid that includes *petRP*, but not in extracts prepared from the strain carrying an empty plasmid (Figure 2E). PetR degradation also required copper supplementation of extracts, demonstrating that PetR degradation depends on this metal (Figure 2E). These results also showed that PetP can degrade PetR directly without the participation of any other protein from *Synechocystis*. Thus, PetP controls PetR levels directly via a copper-dependent mechanism.

### PetR binds to both *petE* and *petJ* promoters

In order to establish whether PetR regulates *petJ* and *petE* directly, we performed Electrophoretic Mobility-Shift assays to analyze PetR binding to both promoter regions. As a first approach, we carried out band shift assays using a recombinant version of PetR (see Materials and Methods for details) and two probes corresponding to the *petJ* and *petE* promoters. PetR was able to bind to both promoters (Figure 3A and B), and the binding was not affected by Cu (Cu^+^ or Cu^2^+; SI Appendix, Figure S8). PetR exhibited a higher affinity toward *petE* than the *petJ* promoter, and was able to completely titrate the probe at lower concentrations (Figure 3A and B). These results indicate a dual role for PetR activating *petJ* expression and repressing *petE*.

**Figure 3.**
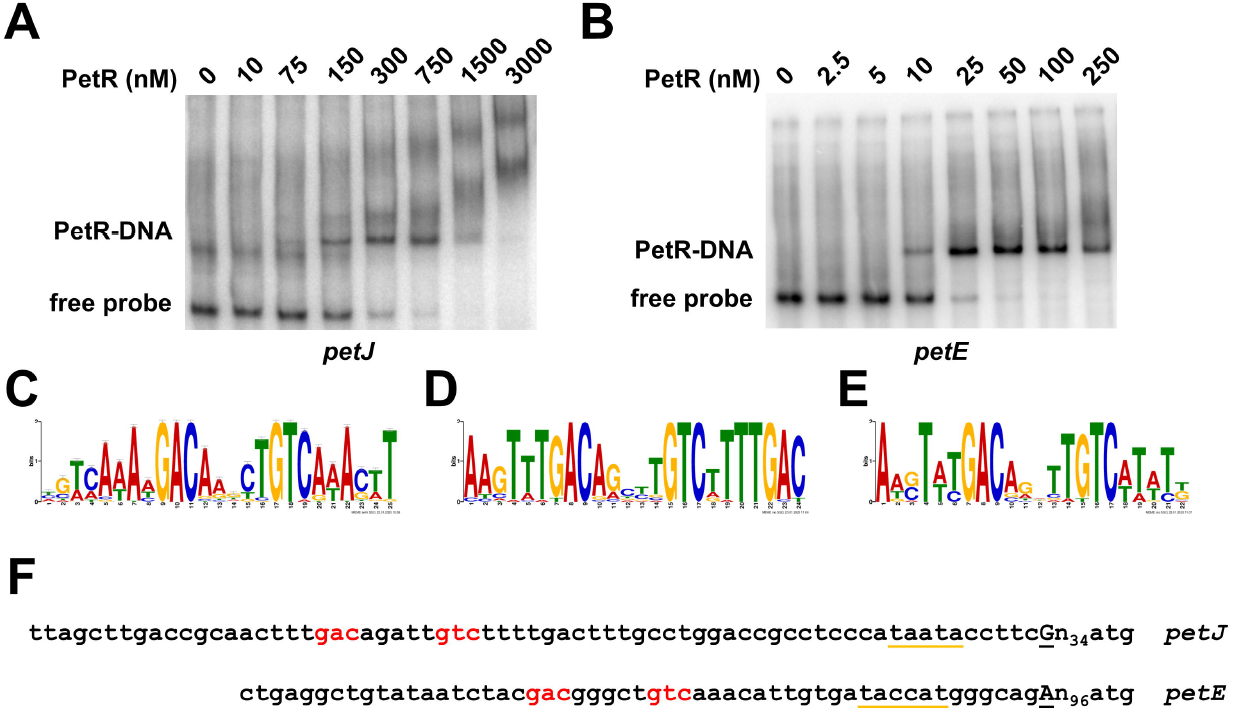
PetR binds to *petJ* and *petE* promoters. A. Electrophoretic mobility shift assay using recombinant PetR to a 103 bp *petJ* promoter probe. The indicated PetR concentration were used in each lane. B. Electrophoretic mobility shift assay using recombinant PetR to a 217 bp *petE* promoter probe. The indicated PetR concentration were used in each lane. C. Sequence logo identified with MEME using upstream sequences (300 bp) from *petE* and *petJ* genes from genomes that also contained *petRP*. D. Sequence logo identified with MEME using upstream sequences (300 bp) from *petJ* genes from genomes that also contained *petRP*. E. Sequence logo identified with MEME using upstream sequences (300 bp) from *petE* from genomes that also contained *petRP*. F. Alignment of the *petJ* and *petE* promoter sequences from *Synechocystis*. Transcriptional start sites are in capital letters and underlined, −10 boxes are yellow underlined and conserved nucleotides from the motif found in C are in red.

In order to identify the PetR binding site, we carried out a bioinformatic analysis of the *petE* and *petJ* promoters in cyanobacterial genomes. A 300-bp fragment surrounding *petE* and *petJ* promoters was retrieved from genomes that contained *petE*, *petJ,* and *petRP* using MGcV database (http://mgcv.cmbi.ru.nl/; SI Appendix, Supplementary dataset S2). Analysis of the sequences of *petE* and *petJ* promoters using MEME suite revealed that the motif shown in Figure 3C was significantly enriched in both. This motif contains the GACN_5_GTC sequence, which was completely conserved, positioned 75 bp upstream from the starting ATG codon (and 37 bp from the tsp) in the *petJ* promoter, and at 118 bp from the ATG codon (and 23 bp from the tsp) in the *petE* promoter. In the *petJ* promoter, the putative binding site is centered around the −35 box in the promoter, in a location suitable as an activator site. In contrast, this sequence is located between the −10 and −35 boxes in the *petE* promoter, compatible with a repressor role of PetR. These locations are consistent with gene expression data obtained by RNA blotting (Figure 1) and transcriptomics analysis (see below) of PETR and PETP mutant strains, which suggests that PetR activates *petJ* and represses *petE* expression. Notably, when MEME analysis was repeated using only *petJ* promoters, an extended motif was identified (Figure 3D). This motif includes the core of the motif shown in Figure 3C, but also two direct repeats of the sequence TTTGAC-N_9_-TTTGAC (Figure 3D). A different motif was also identified when the same analysis was performed using only *petE* promoter sequences (Figure 3E). This motif includes the core motif shown in Figure 3C, but in an extended form of the inverted repeat GACA-N_3_-TGTC. However, the *Synechocystis* sequence in the *petE* promoter is GACG-N_3_-TGTC (Figure 3F).

### RNA-seq analysis of the PetR regulon

To fully characterize the PetR regulon, RNA-seq was carried out using RNA extracted from WT, PETR, and PETP strains grown in BG11C-Cu, and 2 h after addition of 0.5 μM Cu addition (SI Appendix Supplementary dataset S3). Using a threshold of at least a 2-fold change and *P_adj_* < 0.01, 14 genes were induced (*petE,* both plasmid and chromosomal copies of *copMRS, copBAC*, *and nrsBAC*) and three were repressed (*petJ, slr0601, and slr0602*) in the WT strain after copper addition (SI Appendix, Table S1; Supplementary dataset S4), in agreement with previous microarray analysis (27). A similar response was observed in the PETR strain with *copMRS* (both copies)*, copBAC*, and *nrsBAC* operons, and three additional genes of unknown function were also differentially regulated (SI Appendix, Table S1). In the PETP strain, 16 genes were induced after copper treatment, including *copMRS* (both copies)*, copBAC*, and *nrsBAC* operons (SI Appendix, Table S1). As expected from the results shown in Figure 1, neither *petE* nor *petJ* were differentially expressed in PETR or PETP strains (SI Appendix, Figure S9). Furthermore, *slr0601* and *slr0602* were not differentially regulated in PETR or PETP (SI Appendix, Table S1). When gene expression was directly compared between WT and PETR strains, eight genes were differentially regulated in BG11C-Cu (seven expressed at higher levels in WT, and one downregulated), and six were differentially regulated after copper addition (SI Appendix, Supplementary dataset S4). As expected, this group of genes included *petRP* (*petR* was induced 5.93 and 4.73-fold, and *petP* was induced 37 and 38-fold for in BG11C-Cu and +Cu, respectively), *petE* (repressed 41-fold in −Cu and 2-fold in +Cu), and *petJ* (induced 168-fold in −Cu and 8.7-fold in +Cu), as well as *slr0601*, *slr0602*, *slr0242*, *sll1796*, *sll0494*, and *slr1896* (SI Appendix, Supplementary dataset S4). Similarly, when WT and PETP strains were compared, only three genes were differentially regulated in BG11C-Cu (all upregulated), and eight genes were differentially regulated in BG11C+Cu (also upregulated). These genes included *petP* (upregulated 5-fold in both conditions in WT), *petE* (also upregulated in WT, by 5.3-fold in −Cu and 116-fold in +Cu), *petJ* (repressed 25-fold in +Cu), and *slr0242*, *slr0601*, *slr0602*, *slr1896*, and *sll1926* (SI Appendix, Supplementary dataset S4). These results suggest that the PetRP regulon comprises *petJ*, *petE*, *slr0601*, and *slr0602*, as these genes were regulated by copper in WT cells, and affected in both PETR and PETP strains, although with opposite patterns in the two mutant strains.

The *slr0601* and *slr0602* genes are linked in the genome and probably form an operon (SI Appendix, Figure S10A). Expression of *slr0601* was analyzed by RNA blotting, and the pattern was similar to that of *petJ*: repressed after copper addition in WT, not expressed in PETR, and constitutively expressed in PETP strains (SI Appendix, Figure S10B). Furthermore, PetR could bind to the promoter sequence of *slr0601* (SI Appendix, Figure S10C), which contains a sequence matching the one identified in Figure 3 (SI Appendix, Figure S10D). Together, these results indicate that the *petRP* regulon is only composed of *petJ*, *petE*, and *slr0601*-*slr0602* in *Synechocystis*.

### Physiological characterization of PETR and PETP mutant strains

All mutants generated in this study were fully segregated in BG11C, and were able to grow in BG11C-Cu, but given their gene expression programs (Figures 1 and 2; SI Appendix, Figure S9; Table S1, supplementary dataset S4), growth defects were expected in different copper regimes. We analyzed the growth of the PETR and PETP strains on plates differing in copper availability: BG11C-Cu+BCSA (no copper), BG11C-Cu (~30 nM Cu), and BG11C (330 nM Cu). PETR was only affected in BG11C-Cu containing BCSA, similar to the PETJ strain, which has an interrupted *petJ* gene (Figure 4A). By contrast, the PETP strain was not affected under any conditions (Figure 4), because it expressed *petJ* in all tested conditions (Figure 1). Finally, the PETE strain showed a decreased growth rate (but survived) in BG11C, as previously reported (25, 26, 33) (Figure 4A and SI Appendix, Figure S11). A double mutant lacking both genes (PETRP strain) was indistinguishable from the single mutant PETR (Figure 4A and SI Appendix, Figure S6). Analysis of photosynthetic activity revealed that PETJ and PETR strains reached lower oxygen evolution rates (16.8 ± 0.3 μmol min^−1^/OD_750nm_ and 18.9 ± 0.75 μmol min^−1^/OD_750nm_, respectively) than WT (23 ± 3.82 μmol min^−1^/OD_750nm_), PETE (24.5 ± 2.46 μmol min^−1^/OD_750nm_), and PETP (23.2 ± 5 μmol min^−1^/OD_750nm_) strains when cultured in BG11c-Cu+BCSA, although they saturated at the same light intensity (Figure 4B). When cultured in BG11C, the PETE strain exhibited a severe defect in oxygen evolution (reaching only 5.3 ± 2.7 μmol min^−1^/OD_750nm_ and saturating at a light intensity of 180 μmol m^−2^s^−1^(Figure 4C). Surprisingly, the PETP strain exhibited a lower maximum capacity, reaching only 13.24 ± 2.63 μmol min^−1^/OD_750nm_, while WT (24.11 ± 3.66 μmol min^−1^/OD_750nm_), PETJ (22.25±4.34 μmol min^−1^/OD_750nm_), and PETR (21.87 ± 4.36 μmol min^−1^/OD_750nm_) strains all reached similar levels, equivalent to those observed in BG11C-Cu (Figure 4C). Despite this, all strains saturated at the same light intensity (500 μmol m^−2^s^−1^). The PETP strain expressed *petJ* constitutively (Figures 1 and 2), and was therefore not expected to show any photosynthetic defect.

**Figure 4.**
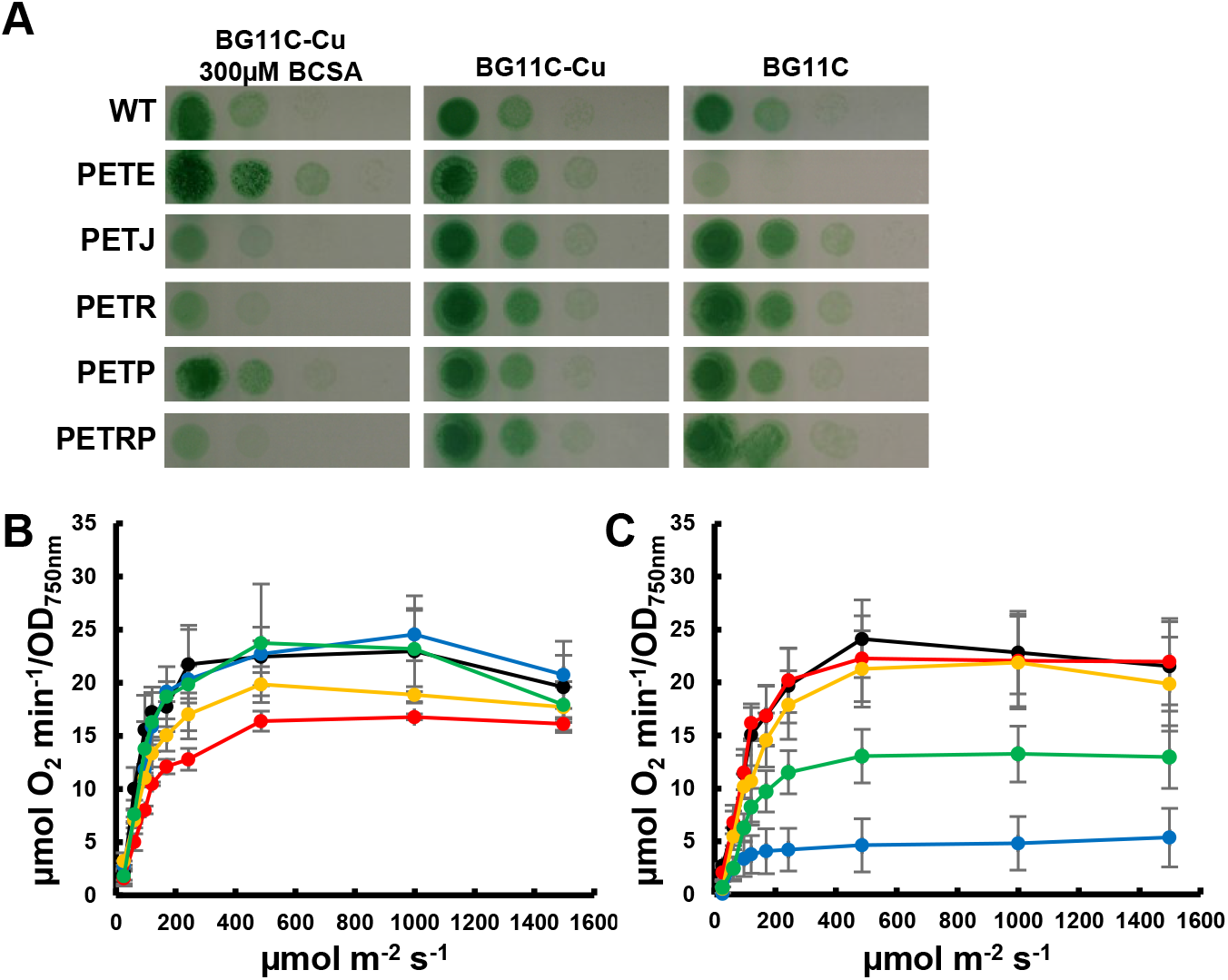
Physiological characterization of *petR* and *petP* mutant strains. A. Growth of WT, PETE, PETJ, PETR, PETP and PETRP strains in different copper availability regimes. Tenfold serial dilutions of a 1 μg chlorophyll mL^−1^ cells suspension were spotted onto BG11C-Cu+BCSA, BG11C-Cu o BG11C. Plates were photographed after 5 days of growth B. Oxygen evolution measured using a Clark electrode at increasing light intensities in exponential growing cultures (OD_750nm_= 0.5-1) of WT 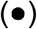, PETJ 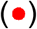, PETE 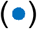, PETR 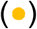 and PETP 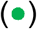 strains grown in BG11C-Cu+BCSA. Data are the mean ± SE of at least three biologically independent experiments. C. Oxygen evolution measured using a Clark electrode at increasing light intensities in exponential growing cultures (OD_750nm_= 0.5-1) of WT 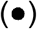, PETJ 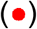, PETE 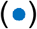, PETR 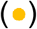 and PETP 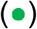 strains grown in BG11C. Data are the mean ± SE of at least three biologically independent experiments.

### Genetic interactions between *petRP*, *petJ*, and *petE*

Our results show that *petRP* is the main system responsible for the *petJ*/*petE* switch following copper addition. Either *petJ* or *petE* are required for photosystem I (PSI) reduction, and therefore for the growth of cyanobacteria (33), despite early studies showing growth of *Synechocystis* in the absence of both proteins (34). The results presented in Figures 1 and 4 suggest that the low levels of PC present in BG11C-Cu (even in the presence of BCSA) were enough to sustain growth of PETR and PETJ mutant strains, albeit at decreased rates (Figure 4A). Furthermore, although the PETE strain showed impaired growth in the presence of copper (Figure 4A), it was able to grow at decreased rates because all cells grew after prolonged incubation (SI Appendix, Figure S11), probably due to low levels of Cyt*c*_6_. Moreover, a double mutant lacking functional *petE* and *petR* genes (or both *petR* and *petP*, strain PETERP) could not be completely segregated (SI Appendix, Figure S12), probably due to low expression levels of *petJ* in the absence of PetR (Figures 1A and SI Appendix, Figure S9), reinforcing the idea that at least one of the soluble transporters (even at low levels) is needed in *Synechocystis*. By contrast, a double mutant lacking *petE* and *petP* (PETEP) was able to grow at WT rates in the presence of copper (SI Appendix, Figure S11 and S12). Consistently, the double mutant PETJP did not segregate, while PETJR and PETJRP mutants fully segregated (SI Appendix, Figure S12). Together, these results suggest that a double mutant lacking both *petJ* and *petE* should not be viable. To test this hypothesis, a double mutant was generated by transforming the PETE mutant with pPETJCm. Merodiploid colonies were obtained, but they never segregated completely (SI Appendix, Figure S13), despite subculturing under several light regimes for more than 2 years. To confirm that PC and Cyt*c*_6_ were expressed in slow-growing mutants, we used concentrated extracts and loaded 150 μg of total protein per lane. Cyt*c*_6_ was detected at low levels in WT and PETE strains, but not in the PETR strain, in the presence of copper (SI Appendix, Figure S12). In WT, PETR, and PETJ strains, PC was detected (although at low levels; Figures 1, 2, and SI Appendix, Figure S12) in the absence of Cu, therefore allowing growth of these strains at decreased rates (Figure 4).

## Discussion

Acquisition of PC as an alternative electron carrier was probably an early adaptation after oxygenic photosynthesis appeared, since local oxygenation (even before the GOE) presumably decreased Fe availability. This is reinforced by the presence of both electron carriers in early-branching cyanobacteria such as *Gloeobacter* and *Pseudoanabaena* (SI Appendix, Figure S1, Supplementary dataset 1). Although known about for more than 40 years (13, 14), the regulatory system responsible for the *petJ*/*petE* switch in cyanobacteria remained undiscovered. Herein, we show that the *petRP* genes are responsible for this regulation (Figures 1 and 5). The system seems to have been acquired early during evolution, because like *petE*, it is present in early branching cyanobacteria, and it is widespread in most cyanobacterial clades (SI Appendix, Figure S1). The only exception is the *Synechococcus*/*Prochlorococcus* clade, in which most strains lack *petRP* genes, similar to previous observations on other regulatory systems (35). We showed that *petRP* act in a single pathway in which PetR positively regulates *petJ* and negatively regulates *petE*, while PetP regulates PetR protein levels in response to copper (Figure 5). PetR directly regulates *petE* and *petJ* gene expression by binding to the promoter regions of these two genes (Figure 3). However, the affinity for both promoters is differs markedly, with the *petJ* promoter requiring much higher PetR concentrations for binding than the *petE* promoter *in vitro*. In addition, binding sites are in different locations in the two promoters, replacing the −35 box in *petJ* and between −10 and 35 in *petE*, consistent with an activating role in *petJ* and a repressing role in *petE*, as confirmed by gene expression analysis (Figures 1, 2 and 5; SI Appendix, Figure S9, Table S1, supplementary dataset S4).

**Figure 5.**
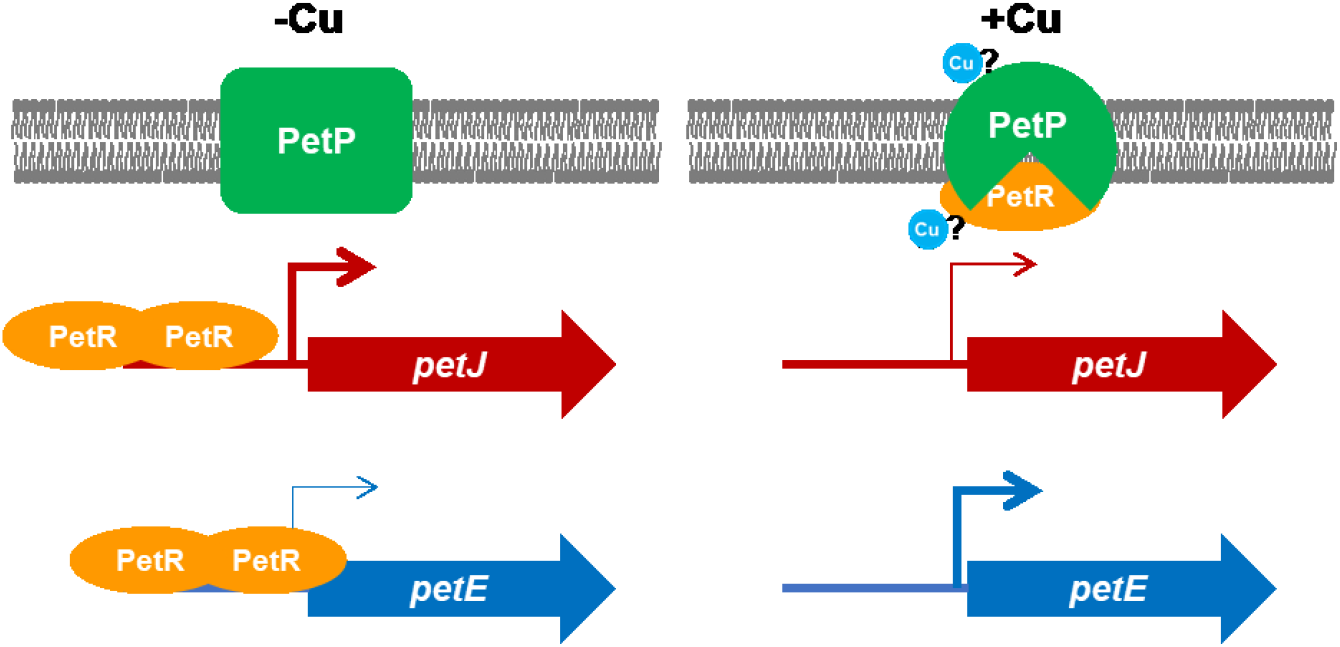
Model of the regulatory mechanism for *petE*/*petJ* switch mediated by PetRP.

Furthermore, our bioinformatics analysis identified an inverted repeat as the core binding site for both promoters in cyanobacteria (Figure 3C), but when *petJ* and *petE* promoters were used, variations of the motif were identified (Figure 3D and E). Given that the protein abundance of PetR *in vivo* depends on copper concentration (Figure 2), and binding to the *petJ* (and *slr0601*) promoter requires higher PetR concentrations (Figure 3 and SI Appendix, Figure S10), *petJ* will only be fully activated when all copper is exhausted (and PetR accumulates at high levels), preventing induction of *petJ* if some copper remains available for PC synthesis. Although BlaI/CopY homologs have been reported to act only as repressors, our results strongly suggest that PetR acts as an activator for *petJ* expression (Figures 1, 3, and SI Appendix, Figure S9B; Table S1). This indicates that BlaI/CopY homologs may also act as activators, as suggested previously (36). Since the binding sites for PetR activation are different to the repressor sites (Figure 3), it is possible that they have remained elusive when studying PetR homologs (37). This dual role for DNA binding proteins that were initially considered to be repressors has been uncovered recently in proteins such as Fur, which was believed to be an activator in both its apo and holo forms (38–40).

Our RNA-seq analysis revealed that only four genes are under the control of the *petRP* system in *Synechocystis*: *petE*, *petJ*, *slr0601*, and *slr0602* (SI Appendix, Figures S9 and S10; Table S1). Inspection of the promoter sequences of these four genes reveals conservation of the core sequence, identified using GACN_5_GTC in the *petJ* and *petE* promoters, but also using only *petJ* promoter sequences (Figure 3 and SI Appendix, Figure S10). The *slr0601* and *slr0602* genes encode two small proteins restricted to cyanobacteria that share low sequence conservation. Slr0601 contains two transmembrane domains with both C-terminal and N-terminal regions predicted to be cytoplasmic, while Slr0602 contains a coiled-coil region but no other recognizable domains. The low levels of sequence conservation and the lack of known domains make it difficult to assign putative functions for these two proteins.

PetP belongs to the M48/M56 family of proteases, and we showed that this protein regulates PetR by controlling its protein abundance in response to copper *in vivo*. This regulation is probably direct as we showed that recombinant PetR is degraded *in vitro* by both *Synechocystis* and *E. coli* extracts, and that this degradation depends on the presence of PetP and copper (Figure 2). This regulatory system is homologous to the BlaI/BlaR regulatory system that regulates β-lactamase expression in *S. aureus*. We explored the mechanism by which copper is sensed by PetRP. PetR belongs to the BlaI/CopY family of transcription factors, and CopY proteins have been demonstrated to bind copper directly (41). Therefore, it is possible that PetR may bind copper, and although this binding may not affect its DNA binding activity (SI Appendix, Figure S8), it may alter its conformation, making it more susceptible to degradation by PetP. Nevertheless, CopY copper binding sites are not conserved in PetR (28), making this hypothesis unlikely.

BlaR/MecR contains a C-terminal extra-cytoplasmic domain that is acylated by β-lactam antibiotics, and this modification activates BlaR, which subsequently degrades BlaI. This domain is absent in PetP, and therefore copper sensing must be mediated by a different domain. The loop between the second and third transmembrane domains of PetP includes two methionine residues (M53 and M58 in *Synechocystis*) that are conserved in all PetP sequences (SI Appendix, Figure S5), which could be putative copper ligands. Interestingly, this domain in BlaR/MecR interacts with the periplasmic C-terminal domain after it binds to β-lactams (42–44). This interaction has been proposed to transduce the signal and activate the protease domain in BlaR. Therefore, these two methionine residues in PetP may bind copper and transduce the signal in a similar way.

Moreover, as PetP is an integral membrane protein, it would be interesting to identify which cyanobacterial membrane system this protein targets. Most copper in *Synechocystis* is presumably sequestered in the thylakoids, where it is bound to PC (45, 46), and CopS, the sensor protein in the copper resistance system, is located to both plasma and thylakoid membranes (25). If PetP is also targeted to both membrane systems, this could allow PetP to respond not only to an external surplus of copper, but also when the metal is released from internal PC. PetP localization to the thylakoids is reinforced by the fact that mutants lacking the P_1_-type-Cu-ATPases CtaA, PacS, or both (which are involved in Cu acquisition by PC) displayed decreased levels of *petE* mRNA after copper addition (25, 47). This suggests that Cu needs to be transported to the thylakoid in order to be detected by the PetRP system.

Furthermore, our results showed that *Synechocystis* can synthesize PC even when copper levels are very low due to the presence of BCSA in BG11C-Cu. This indicates that some copper can be transported even in the presence of BCSA, suggesting that the copper import system (which remains unidentified) has a higher affinity for copper than the chelating agent. Accumulation of PC in the presence of BCSA was only observed in the PETR strain in which *petE* expression was constitutive (Figures 1, 5, and SI Appendix, Figure S9), because in the WT strain *petE* is repressed and PC cannot be synthesized, even if copper is available.

Finally, our results also suggest that there are other layers of regulation at the posttranscriptional level because Cyt*c*_6_ and PC protein abundance does not always correlate with mRNA levels. PC was more abundant in Cu-containing media in the PETR strain in which *petE* mRNA levels did not change in response to copper (Figure 1). Similarly, Cyt*c*_6_ levels were decreased in the PETP strain after copper addition, even when mRNA levels were not decreased. PC protein abundance may be related to the protein not being correctly translated and/or folded in the absence of available copper, or due to degradation of the apoprotein, but synthesis of Cyt*c*_6_ should not be altered by copper. It has been proposed that IsaR sRNA regulates *petJ* translation in response to Fe deprivation (24). We did not find *isaR* to be differentially regulated in our RNA-seq experiment (SI Appendix, Figure S14), hence it is unlikely that this regulation is mediated by *petRP*. Furthermore, oxygen evolution was affected in the PETP strain (which expresses Cyt*c*_6_; Figure 4C) in the presence of copper, suggesting that additional changes in the photosynthetic transport chain might be present, as reported for *Chlamydomonas* (48). Finally, although single mutants of either *petJ* or *petE* were able to grow, even in a copper regime in which expression of the other gene was repressed, these strains exhibited growth and photosynthesis defects (Figure 4), in agreement with previous studies (33, 49–51). Double mutants of both *petJ* and *petE* were not viable (SI Appendix, Figure S13), and double mutants lacking one of the genes encoding the electron transport proteins and with decreased expression of the other due to the lack of *petP* (constitutive repression of *petE;* strain PETJP) or *petR* (unable to induce *petJ*; strain PETER) were not viable either (SI Appendix, Figure S12). Together, these results demonstrate that *petRP* are essential for expression of these genes, and that levels of PC or Cyt*c*_6_ proteins must be above a critical threshold for survival of the strains. These results also rule out the existence of another electron transfer mechanism between cytochrome *b*_6_*f* and PSI.

## Materials and Methods

### Strains and culture conditions

All *Synechocystis* strains used in this work were grown photoautotrophically on BG11C (52) or BG11C-Cu (lacking CuSO_4_) medium at 30°C under continuous illumination using 4000–4500 K white-LED light (50 μmol photon m^−2^ s^−1^, measured at the flask surface) and bubbled with a stream of 1% (v/v) CO_2_ in air. BG11C-Cu was supplemented with 300 μM BCSA as a chelating agent to trap any trace of copper, when required. For plate cultures, media were supplemented with 1% (w/v) agar. Kanamycin, nourseothricin, chloramphenicol, spectinomycin, and erythromycin were added to a final concentration of 50 μg mL^−1^, 50 μg mL^−1^, 20 μg mL^−1^, 5 μg mL^−1^, and 5 μg mL^−1^, respectively. Experiments were performed using cultures from the logarithmic phase (3–4 μg chlorophyll mL^−1^). *Synechocystis* strains are described in SI Appendix, Table S2.

*E. coli* DH5α cells were grown in Luria Broth or M9 (supplemented with 1 mL LB per L of M9) media and supplemented with 100 μg mL^−1^ ampicillin, 50 μg mL^−1^ kanamycin, 50 μg mL^−1^ nourseothricin, 20 μg mL^−1^ chloramphenicol, and 100 μg mL^−1^ spectinomycin when required.

### Insertional mutagenesis of Synechocystis genes

For *slr0240* (*petR*) mutant construction, a 1524-bp fragment was amplified using oligonucleotides 223 and 224, cloned into pSPARKII (generating pPETR), and a Sp^R^ cassette was inserted in the *Eco*RV site at codon 31 in the *slr0240* open reading frame (ORF), generating plasmid pPETR_Sp. For the *slr0241* (*petP*) mutant, a 970-bp fragment was amplified using oligonucleotides 248 and 249 and cloned into pSPARKII, generating pPETP. A 582-bp *Nhe*I fragment was substituted by an *Xba*I erythromycin cassette, generating pPETP_Ery. For the *petRP* double mutant, a 1492-bp fragment was amplified by PCR using oligonucleotides 256 and 257, and cloned into the pSPARKII vector, generating the pPETRP plasmid. A 650-bp *Bst*EII fragment from plasmid pPETRP was substituted by an *Hin*cII-digested SpR cassette from pRLSpec, generating pPETRP_Sp. To generate the *petJ* mutant, a 1023-bp fragment was amplified using oligos 235 and 236, and cloned into pSPARKII, generating pPETJ, and a *Xba*I CC1 Cm resistance cassette was inserted in the *Nhe*I site present in the *petJ* ORF, generating pPETJ_Cm. For PETR complementation, a 639-bp fragment (185 bp upstream of the *petR* STOP codon) was amplified using oligonucleotides 256 and 234, and cloned into pGLNN digested with *EcoR*I and *Not*I, generating pPETR_comp. For the PETR3 strain, a 463-bp fragment was amplified using oligonucleotides 258 and 234 (adding *Sal*I and the pET28 ribosome binding site upstream of *petR* ATG), digested with *Sal*I-*Not*I, and cloned into pGLN-PcpcB digested in the same way, generating pPETR_OE. These plasmids were used to transform WT *Synechocystis*, PETR, PETE, or PETJ strains, generating PETR, PETR2, PETR3, PETP, PETRP, PETER, PETEP, PETERP, PETJR, PETJP, PETJRP, and PETEJ strains. Segregation of the strains was verified by PCR with the appropriate primer pairs, as depicted in SI Appendix, Figures S2, S3, S11, and S13. All plasmids were sequenced to verify that no unwanted mutations were introduced. Sequences for all oligonucleotides are listed in SI Appendix, Table S3.

### RNA isolation and RNA blot analysis

Total RNA was isolated from 30 mL samples of *Synechocystis* cultures in the mid-exponential growth phase (3 to 4 μg chlorophyll mL^−1^). Extractions were performed by vortexing cells in the presence of phenol-chloroform and acid-washed baked glass beads (0.25–0.3 mm diameter) as previously described (53). A 5 μg sample of total RNA was loaded onto each lane of a 1.2% denaturing agarose formaldehyde gel (54), electrophoresed, and transferred to a nylon membrane (Hybond N-Plus; GE Biosciences). Prehybridization, hybridization, and washes were performed in accordance with the manufacturer’s instructions. Probes for RNA blot hybridization were synthesized by PCR using oligonucleotide pairs 172-173, 174-175, 389-390, 275-276, 274-275, and 178-179 (SI Appendix, Table S3) for *petE*, *petJ*, *copM*, *slr0601*, and *rnpB*, respectively. DNA probes were ^32^P-labeled with a random primer kit (Rediprime II catalog #RPN1633; GE Biosciences) using [α-32P] dCTP (3,000 Ci/mmol). Hybridization signals were quantified with a Cyclone Phosphor System (Packard). Each experiment included at least three biological replicates.

### Immunoblotting

For analysis, proteins were fractionated by SDS-PAGE and immunoblotted with antibodies against plastocyanin (1:3000), cytochrome *c*_6_ (1:3000), CopM (1:7000), PetR (1:6000), or *Synechococcus* sp. PCC 6301 glutamine synthetase I (1:100,000). ECL Prime (catalog # RPN2232, GE Biosciences) was used to detect the different antigens with anti-rabbit secondary antibodies conjugated to horseradish peroxidase (1:25,000; catalog # A0545, Sigma). Either photographic film or a ChemiDoc Imaging System (Bio-Rad) were used for signal detection and quantification.

Samples were prepared from whole cells or soluble extracts. For whole cells, cells were harvested from cultures with an optical density at 750 nm (OD_750nm_) of 1, the supernatant was carefully removed, cells were resuspended in 100 μL of 1× Laemmli loading buffer, and boiled for 10 min. For soluble extracts, Cells equivalent to 20 OD_750nm_ were collected, resuspended in 300 μL of buffer A (50 mM TRIS HCl pH 8, 50 mM NaCl, 1 mM phenylmethylsulfonyl fluoride) and broken using glass beads via two cycles, separated by 5 min on ice, of 1 min in a minibead beater. Cell extracts were recovered from the beads and samples were clarified by two sequential centrifugations: 5 min at 5000 *g* to eliminate cell debris, and 15 min at 15,000 *g* to remove membranes. Protein concentrations in cell-free extracts and purified protein preparations were quantified using Bradford reagent (catalog # 5000006, Bio-Rad), with ovalbumin as a standard, and specified amounts of protein were separated by SDS-PAGE.

### PetR protein purification

The complete *petR* ORF was cloned from *Synechocystis* DNA after PCR amplification with oligonucleotides 233 and 234, and cloned into the *Bam*HI-*Not*I sites of pGEX6P, generating pGEX_PETR. The GST-PetR fusion protein was expressed in *E. coli* DH5α cells. A 1 L culture was grown in Luria broth medium to an OD_600nm_ of 0.6, cooled to 4°C, 0.1 mM isopropyl-b-D-thiogalactopyranoside (IPTG) was added, and culturing was continued at 25°C overnight. Cells were harvested by centrifugation, resuspended in 5 mL of PBS buffer (150 mM NaCl, 16 mM Na_2_HPO_4_, 4 mM NaH_2_PO_4_, 4 mM phenylmethylsulfonyl fluoride, 7 mM β-mercaptoethanol) supplemented with 0.1% Triton X-100, broken by sonication on ice, and insoluble debris was pelleted by centrifugation for 45 min at 25,000 *g*. Soluble extracts were mixed with 1 mL of glutathione agarose beads (catalogue number # 17075601, GE Healthcare) and incubated for 2 h at 4°C with gentle agitation. Beads were transferred to a column and washed extensively with PBS buffer (20–30 column volumes) until no more protein was eluted from the column. The column was equilibrated with 10 volumes of 50 mM TRIS HCl pH 7.5, 150 mM NaCl, 1 mM EDTA, and 1 mM dithiothreitol, and beads were resuspended in two volumes of the same buffer. A 20 μL sample of Precission protease (2 U/μL; catalogue # 27084301, GE Healthcare) was added and beads were incubated overnight at 4°C. Beads were poured into an empty column and the flow through fraction was collected. This fraction was diluted three times in 50 mM TRS HCl pH 8, and applied to a HiTrap Heparin column (catalogue # 17040601, GE Healthcare) connected to an Amersham FPLC system. The column was washed with 10 volumes of 50 mM TRIS HCl, 100 mM NaCl, and a 20 mL gradient from 0.1–1 M NaCl was applied. Fractions containing PetR were pooled and concentrated. GST-PetR fusion protein was eluted from glutathione agarose beads with 3 mL of 50 mM TRIS HCl pH 8 containing 10 mM reduced glutathione after washing the column with PBS. A gel of the purified fractions is shown in SI Appendix, Figure S15.

### Protease assays

Cells equivalent to 20 OD_750nm_ of *Synechocystis* strains were harvested from mid-exponential growth phase cultures, resuspended in 300 μL of buffer containing 50 mM TRIS-HCl pH 8, 50 mM NaCl, 10% glycerol, and 1 mM ZnSO_4_, and broken with 2 volumes of glass beads using a minibead beater. Total cell extracts were collected by piercing the bottom of the tube with a needle. Extracts were centrifuged for 5 min at 5000 *g* to remove the remaining glass beads and unbroken cells, and the supernatant constituted the total extract. A 2-μg sample of GST-PetR was added to 150 μL of total extracts (containing 150 μg of total protein) and incubated at 30°C. At the indicated times, 20 μL samples were removed, mixed with 7 μL of 4× Laemmli buffer and boiled for 10 min. A 5 μL sample was used for immunoblotting using α-PetR antibodies.

For *E. coli* assays, DH5α cells transformed with pN_petRP_Nat or pN_Nat were grown to late exponential phase (OD_595nm_ 1–2), and cells at an OD_595nm_ value of 20 were collected and processed as described above for *Synechocystis*. The pN_petRP_Nat plasmid was constructed by excising a 1478-bp fragment from pPETRP using *Eco*RI and *Xho*I, containing petRP and the 185-bp region upstream from *petR* ATG, and inserting it into pN_Nat (30) cut in the same way.

### Band shift and gel retardation assays

Probes were PCR-synthesized using oligonucleotides 53 and 92 for the *petE* promoter (217 bp), 299 and 262 for the *petJ* promoter (103 bp), and 295 and 296 for *slr0601* (177 bp), introducing a *Sal*I restriction site in all cases. The resulting DNA was digested with *Sal*I and end-labeled with [α-32P]-dCTP (3000 Ci mmol^−1^) using Sequenase v2.0 (Product #70775Y, Affymetrix). The binding reaction was carried out in a final volume of 20 μL containing 4 ng of labeled DNA and 4 μg of salmon sperm DNA in 20 mM TRIS-HCl (pH 8.0), 150 mM KCl, 10 mM DTT, 1 mM EDTA, 10% glycerol, and different amounts (from 0.001 μg to 1 μg) of purified PetR. The mixtures were incubated for 30 min at room temperature and loaded on a Tris/Borate/EDTA (TBE) buffer non-denaturing 6% polyacrylamide gel (acrylamide:bisacrylamide 30:0.6). Electrophoresis was carried out at 4°C and 200 V in 0.25× TBE. Gels were transferred to a Whatman 3 MM paper, dried, and autoradiographed using a Cyclone Phosphor System (Packard). Each experiment was performed at least three times with two independent PetR preparations.

### RNA-seq

The quantity and quality of total RNA were evaluated using RNA electropherograms acquired by an Agilent 2100 Bioanalyzer; Agilent Technologies, Santa Clara, CA, USA), and library construction of cDNA molecules was carried out using Illumina Kapa Stranded Total RNA with a Ribo-Zero Library Preparation Kit. The resulting DNA fragments (DNA library) were sequenced on an lllumina HiSeq 4000 platform using 150-bp paired-end sequencing reads. Sequencing was carried out by STAB VIDA (Lda, Lisbon, Portugal). Reads were aligned against the NCBI genome sequence for *Synechocystis* (NC_000911.1, NC_005229.1, NC_005230.1, NC_005231.1, and NC_005232.1) using bowtie2 version 2.2.4 (55). Raw read counts were calculated using the HTSeq-count function of HTSeq framework 0.11.0 (56). The Bioconductor DESeq2 R software package (57) was used to detect differentially expressed genes. An adjusted *p*-value of <0.01 was considered significant. For RNA-seq density profiles, data were normalized using the BamCoverage function of DeepTools2 0.8.0 (58), and visualized with Integrative Genomics Viewer (IGV) 2.3.6 (59). The RNA-seq dataset is available at GEO accession GSE155385.

### Phylogenomic analysis

16S RNA from 169 cyanobacterial genomes (Supplementary Material S1; all completed genomes from NCBI and those included in (60)) were aligned with MUSCLE 3.8.31, and a maximum likelihood (ML) phylogeny was estimated with IQ-TREE 1.6.11 (61) using the best-fitting model (automatically selected by the ModelFinder function) and selecting the phylogeny with the highest likelihood from those found among independent searches, with 1000 bootstrap replicates. *petE, petJ, petR*, and *petP* genes were searched using TBLASTN. Only hits with ≥50% query coverage and >25% identity were considered homologues, and checked manually. Phylogenetic trees were displayed and edited using the Interactive Tree Of Life webtool iTOL v4 (62).

### Oxygen evolution

Oxygen evolution was measured by a Clark-type oxygen electrode (Hansatech Chlorolab 2) using mid-logarithmic (OD_750nm_ = 0.8–1) cultures adjusted to OD_750nm_ = 0.5 in BG11C or BG11C-Cu+BCSA media supplemented with 20 mM NaHCO_3_ and white LED light.

## Supporting information

Supplementary data set 1

Supplementary data set 2

Supplementary data set 3

Supplementary data set 4

## Acknowledgments

We thank Jose Luis Crespo, Manuel J. Mallén, Miguel Roldán, and Sandra Díaz-Troya for critical reading of the manuscript. This research was funded by grant numbers BIO2016-75634-P and PID2019-104513GB-I00 from the Ministerio de Economía y Competividad (MINECO) and the Agencia Estatal de Investigación (AEI), respectively, and by Junta de Andalucía Group BIO-284, co-financed by European Regional Funds (FEDER), to Francisco J. Florencio. RMGC is the recipient of a predoctoral contract from Ministerio de Educación, Cultura y Deporte, Spain (contract #FPU2015/05025).

