## Supplementary tables for "A protease-mediated mechanism regulates the cytochrome *c*_6_/ plastocyanin switch in *Synechocystis* sp. PCC 6803"

**A copper activated protease regulates the cytochrome $c_6$/ plastocyanin switch in cyanobacteria**

.

1. Instituto de Bioquímica Vegetal y Fotosíntesis, Universidad de Sevilla- CSIC, c/Américo Vespucio 49, 41092 Sevilla, Spain.

2. Departamento de Bioquímica Vegetal y Biología Molecular, Facultad de Biología, Universidad de Sevilla, Avenida Reina Mercedes s/n, 41012 Sevilla, Spain

3. Present address: Centro de Biotecnología y Genómica de Plantas UPM – INIA Parque Científico y Tecnológico de la U.P.M. Campus de Montegancedo, Autopista M-40, Km 38, Pozuelo de Alarcón, Madrid 28223, Spain

* Luis López-Maury

**This PDF file includes:**

Figures S1 to S15
Tables S1 to S3
Legends for Datasets S1 to S4

**Other supplementary materials for this manuscript include the following:**

Datasets S1 to S4

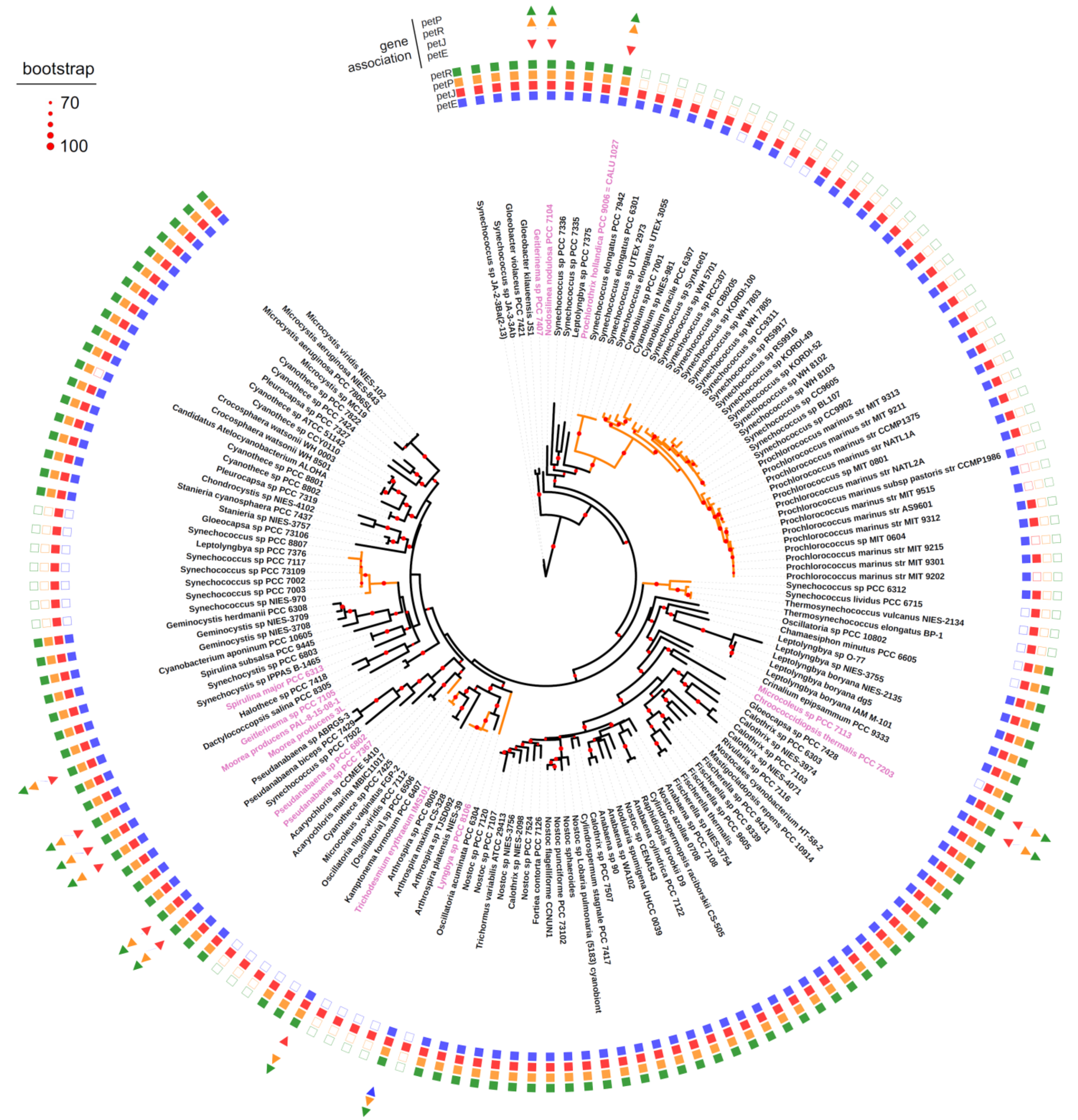

**Figure S1. Conservation of *petRP*, *petE* and *petJ* genes in genomes of cyanobacteria**
Maximum Likelihood phylogenetic tree of cyanobacteria genomes. The tree was estimated using the 16S RNA sequences from 169 cyanobacteria genomes (see Material and Methods section) with IQ-TREE using the best fitting model. Nodal support values are represented with red circles as shown in legend, size correspond to ultrafast bootstrap values from 70 to 100, as implemented in IQ-TREE. Presence of *petE, petJ, petR* and *petP* genes is indicated with filled squares. Genomes that show association of *petE* or *petJ* with *petRP* are highlighted in pink and gene order is also shown by arrows. Orange clades indicate absence of both *petR* and *petP genes*.

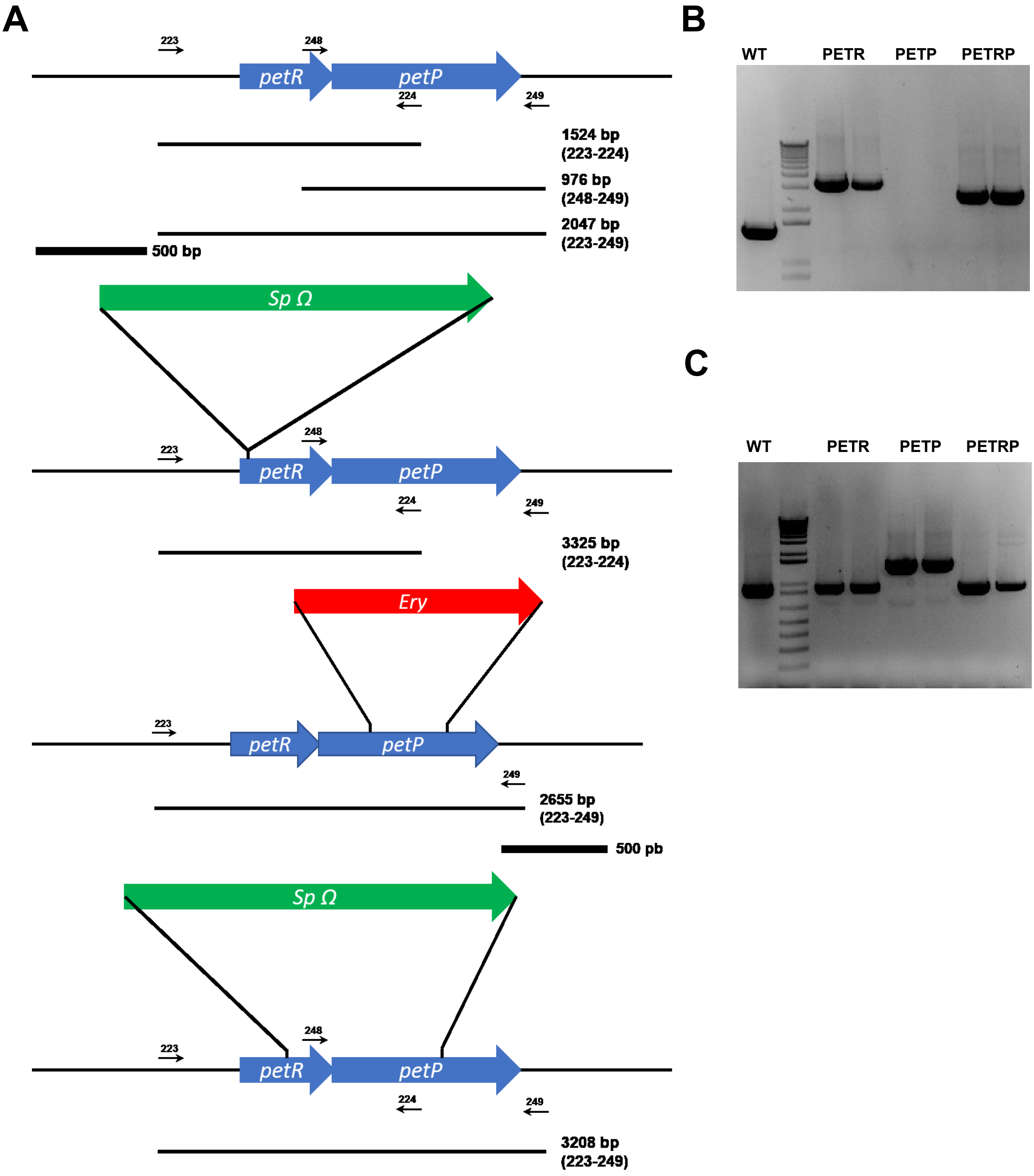


**Figure S2. Construction of PETR, PETP and PETRP mutant strains.**

A. Schematic representation of the *petRP* (*slr0240-41*) genomic region in WT, PETR, PETP and PETRP mutant strains. Oligonucleotides and sizes of bands amplified by the different oligo pairs in all strains are also shown
B. PCR analysis of the *petRP* locus in WT, PETR, PETP and PETRP strains (two clones were analysed for each strain) using oligonucleotides 223-224.
C. PCR analysis of the *petRP* locus in WT, PETR, PETP and PETRP strains (two clones were analysed for each strain) using oligonucleotides 248-249.

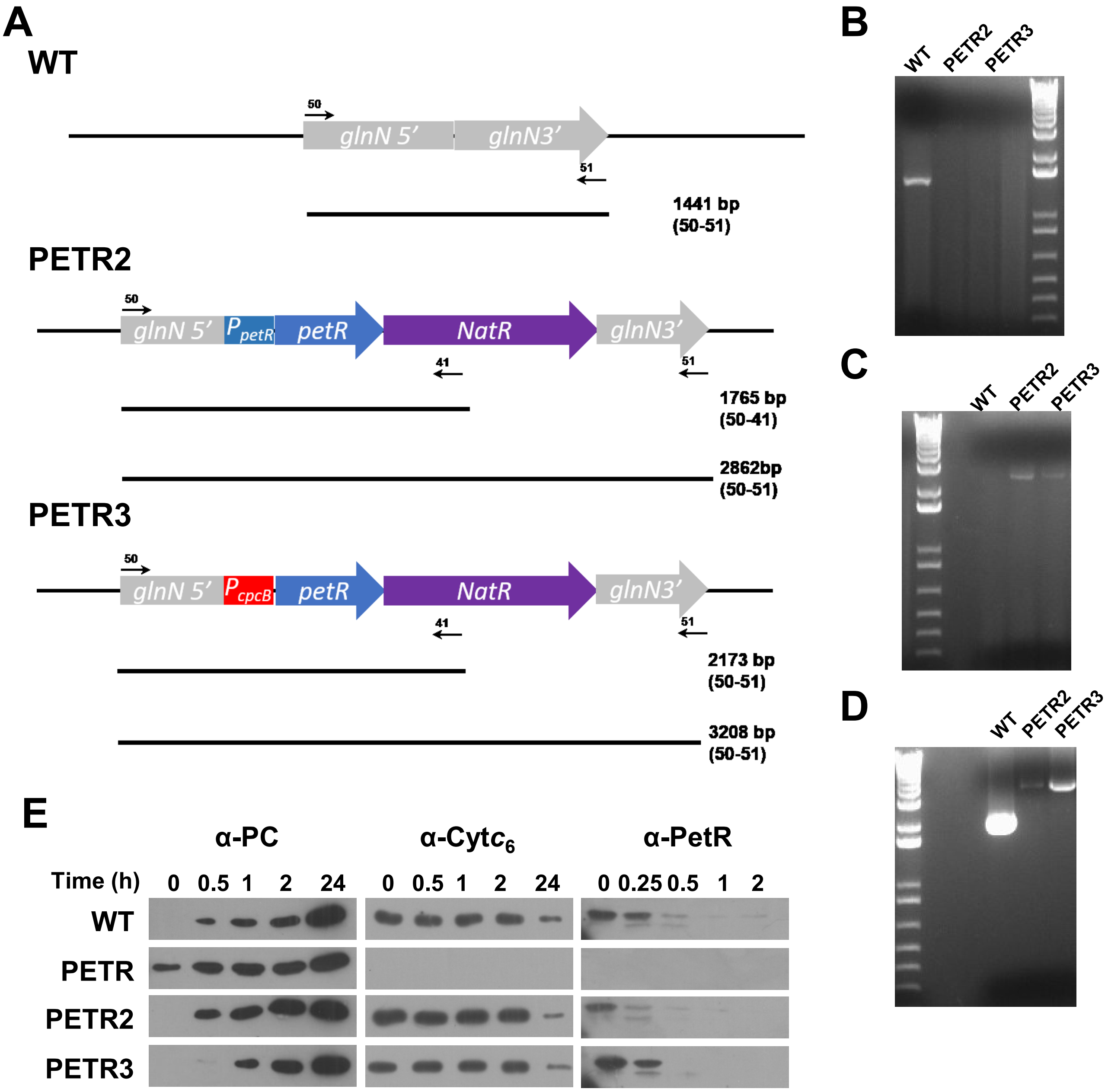


**Figure S3. Complementation of PETR mutant strain.**

A. Schematic representation of the *glnN* genomic region in WT, PETR2 and PETR3 mutant strains. Oligonucleotides and sizes of bands amplified by the different oligo pairs in all strains are also shown
B. PCR analysis of the *petRP* locus in WT, PETR2 and PETR3 mutant using oligonucleotides 223-224. For expected band sizes are depicted in Figure S2
C. PCR analysis of the *glnN* locus WT, PETR2 and PETR3 mutant strains using oligonucleotides 50-51.
D. PCR analysis of the *glnN* locus WT, PETR2 and PETR3 mutant strains using oligonucleotides 50-41.
E. Immunoblot analysis of Cyt$c_6$, PC, and PetR in WT, PETR, PETR2 and PETR3 strains in response to 0.5 μM copper addition. Cells were grown in BG11C-Cu medium and cells were harvested at the indicated times after addition of copper 0.5 μM. 25 μg of total protein from soluble extracts was separated by 15 % SDS-PAGE and subjected to inmunoblot to detect PC, Cyt$c_6$ and PetR. For PetR whole cell extracts were used and 20 μl (equivalent to 0.2 $OD_{750nm}$) were loaded per lane.

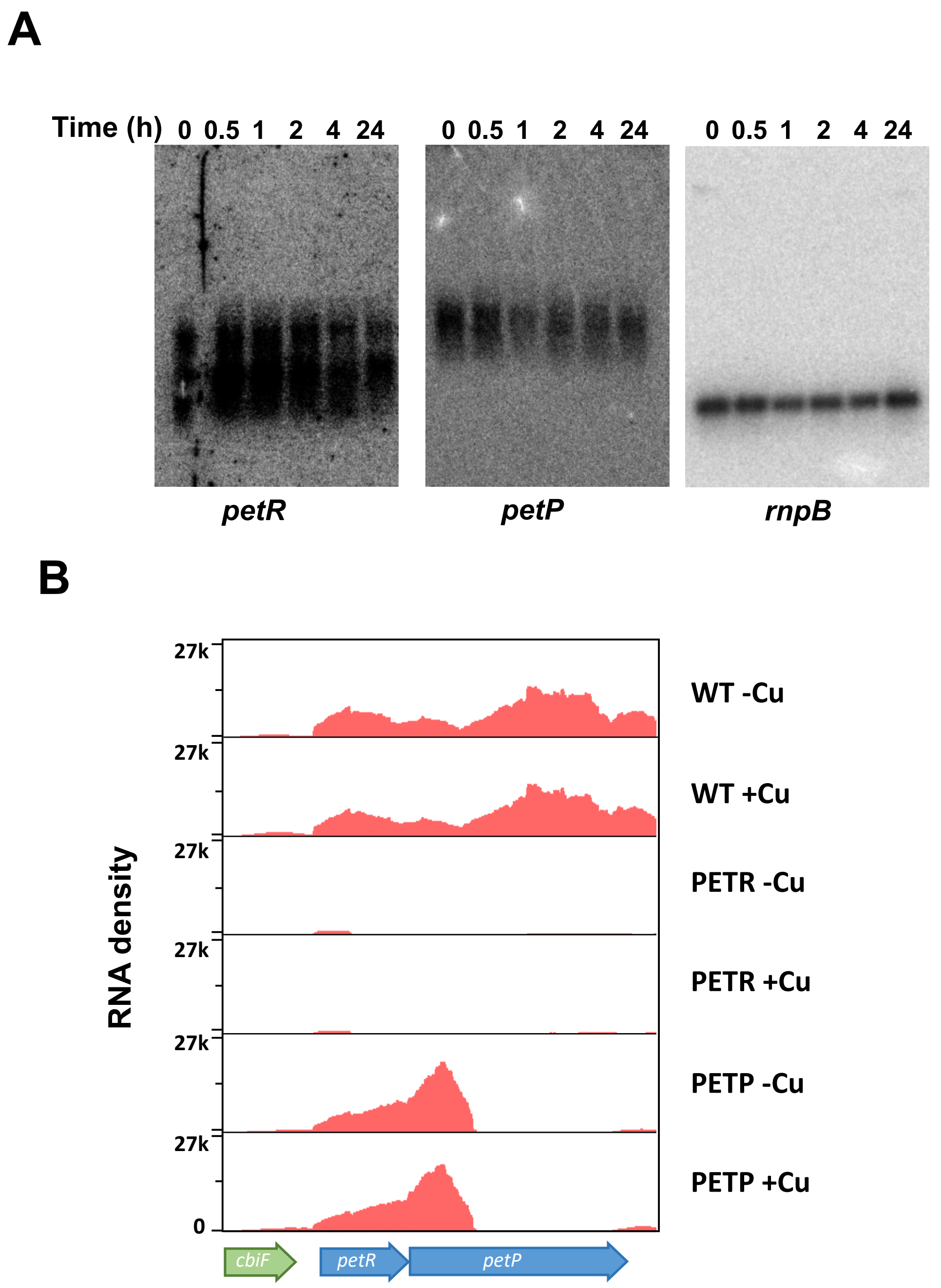


**Figure S4. *petRP* are expressed as a single transcriptional unit.**

A. RNA blot analysis of *petR* and *petP* in the WT strain in response to 0.5 μM copper addition. Total RNA was isolated from cells grown in BG11C-Cu medium at the indicated times after addition of 0.5 μM of copper. The filters were hybridized with *petR* and *petP* probes and subsequently stripped and re-hybridized with a *rnpB* probe as a control.

B. RNA seq density profile of *petR* and *petP* genes in WT, PETR, PETP strains grown in BG11C-Cu medium and after copper treatment.

A

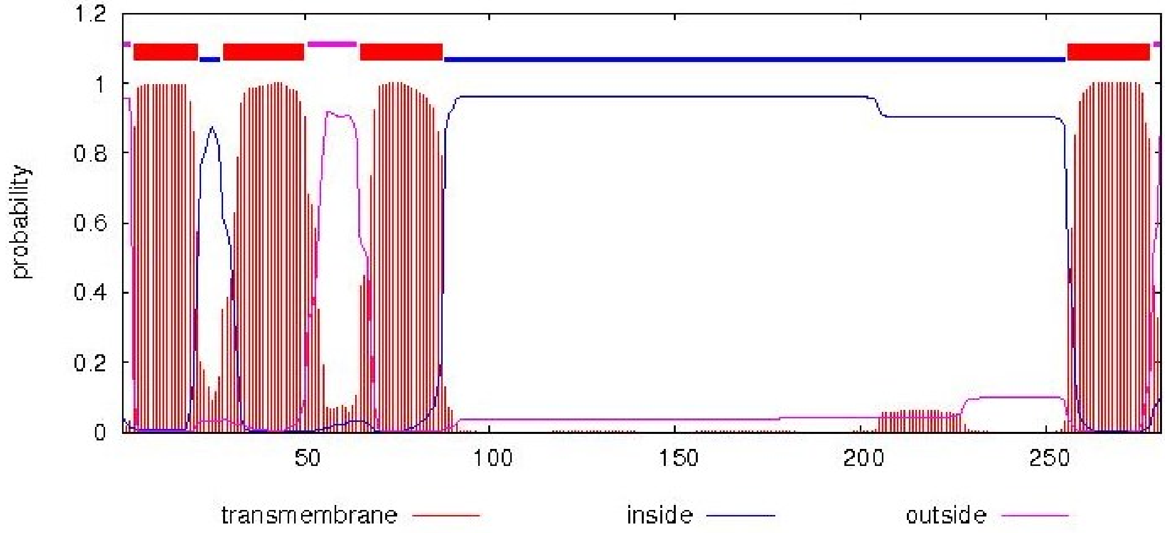


B

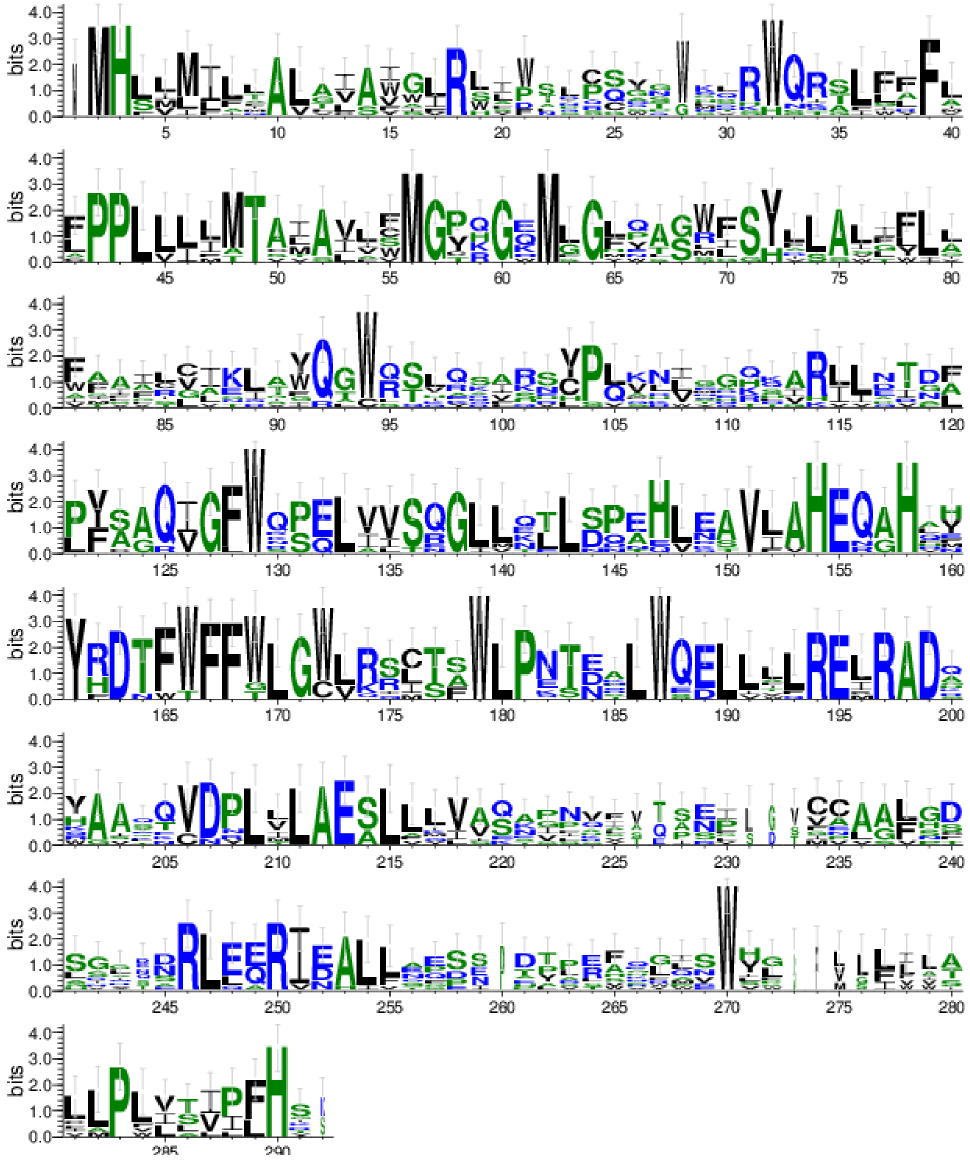


**Figure S5. In silico analysis of PetP sequence.**

A. Prediction of the transmembrane domains in *Synechocystis'* PetP sequence using TMHMM Server v. 2.0.

B. Sequence logo of PetP proteins form cyanobacteria. PetP proteins were identified using BLAST at NCBI (269 sequences). Protein sequences were aligned with MUSCLE using the default parameters and the alignment was submitted to weblogo3 (http://weblogo.threeplusone.com/ ) to generate the consensus sequence shown.

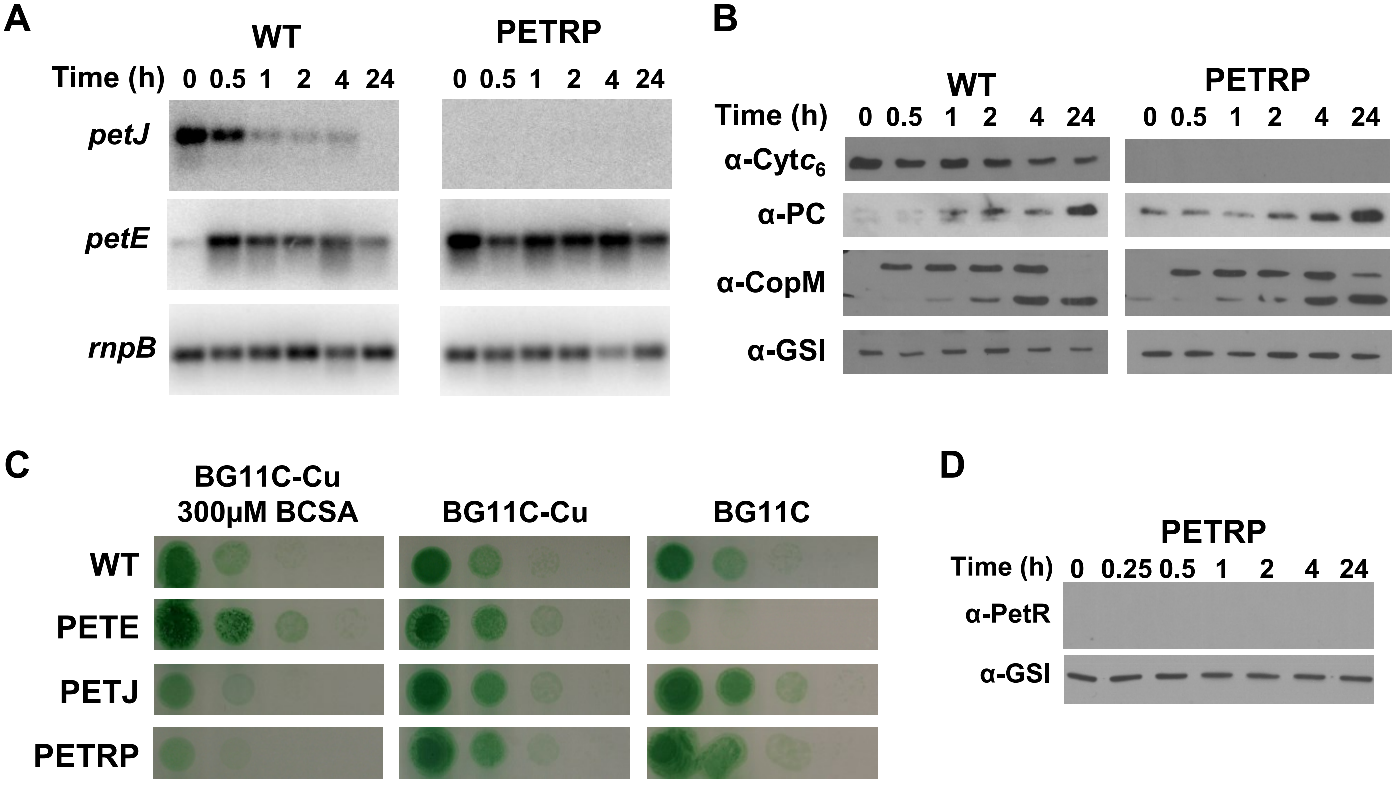


**Figure S6. PETRP strain is indistinguishable from a PETR mutant.**

A. RNA blot analysis of *petE*, *petJ* and *copM* in WT, and PETRP strains in response to 0.5 μM copper addition. Total RNA was isolated from cells grown in BG11C-Cu medium at the indicated times after addition of 0.5 μM of copper. The filters were hybridized with *petJ* and *petE* probes and subsequently stripped and re-hybridized with a *rnpB* probe as a control.
B. Immunoblot analysis of PC, Cyt$c_6$, CopM and GSI in WT and PETRP strains in response to 0.5 μM copper addition. Cells were grown in BG11C-Cu medium and harvested at the indicated times after addition of 0.5 μM of copper. 10 μg of total protein from soluble extracts were separated by 15 % SDS-PAGE and subjected to western blot to detect PC, Cyt$c_6$, CopM or GSI.
C. Growth of WT, PETE, PETJ, PETR, PETP and PETRP strains in different copper availability regimes. Ten-fold serial dilutions of a 1 μg chlorophyll $mL^{-1}$ cells suspension were spotted onto BG11C-Cu+BCSA, BG11C-Cu o BG11C. Plates were photographed after 5 days of growth.
D. Immunoblot analysis of PetR and GSI in PETRP strain grown BG11c-Cu after addition of 0.5 μM copper. Whole cells were loaded (0.2 $OD_{750nm}$), separated by 15 % SDS-PAGE and subjected to immunoblot to detect PetR or GSI.

```
Leptolyngbya_sp_PCC_7375           MSKFPAYRPKRLSLGPLETEILDILWQLGTTSATAIHNQILEDIDRDLTYSSVATVLRRL
Pseudoanabaena_sp_PCC_6802         MVPLPNYRPKQLSLGPLETEILHLIWELGTTTAREIHDRILSDPDRELTYSSVITVLSRL
Synechocystis                      MSWIPPYRPQQLSLGPLEQEILQIIWQLGQATVKDIHDRILSDPDRELAYSSVTTVLNRL
Synechococcus_sp_JA-2-3B'a(2-13)   --MLPKHRPKQLSLGPLESEILDILWDLGSASTRQIHERILADPDRELAYASVTTVLQRL
Acaryochloris_marina_MBIC11017     MASLPDFRPKQLSLGPLEFEILNLIWDLGTATVKQVHEQILTNPDRELAYTSVTTVLNRL
Anabaena7120                       MAPLPDYRPKQMSVGPLEAEILNIVWEVGSATVKDVHDRILADPNRELAYTSVTTVLRRL
Bacillus_licheniformis_BlaI        MKKIP-------QISDAELEVMKVIWKHSSINTNEVIKEL--SKTSTWSPKTIQTMLLRL
Staphylococcus_MecI                ------MDNKTYEISSAEWEVMNIIWMKKYASANNIIEEI--QMQKDWSPKTIRTLITRL
Staphylococcus_BlaI                ------MANKQVEISMAEWDVMNIIWDKKSVSANEIVVEI--QKYKEVSDKTIRTLITRL
                                               .:.  * ::: ::*     ..  :   :  .     :  :: *:: **

Leptolyngbya_sp_PCC_7375           EKKGWIAKRR----VGRAY--HWEPLLSANNAKILESHERLHQFLAASNPDIVAAFADTL
Pseudoanabaena_sp_PCC_6802         VKKGWLTSHK----RGKIL--FWQAAISRPEAQALEAHERLNRFLEVGNADIVAAFADEL
Synechocystis                      TKKGWLVCHR----QGKAF--IWTARVSADQAKAVQSYEQLQQFLAISNPDVVAAFADSL
Synechococcus_sp_JA-2-3B'a(2-13)   SQKGWVSCERRTDPQGRNLPMLWRPRLSRQEAASLRAFTHLQQFLAVGDPDTVAAFADSL
Acaryochloris_marina_MBIC11017     TKKGWLACDK----QNRSF--VWRPLVSRAQANSLWAYDQLQQFLAVGNPDTVAAFADHL
Anabaena7120                       TDKGWLACDK----KGRAF--YWRPLLSKQQAQVIKAHDQLHSFLAVGNPDVVAAFADSL
Bacillus_licheniformis_BlaI        IKKGALNHHK----EGRVF--VYTPNIDESDYIEVKSHSFLNRFYNGTLNSMVLNFLEN-
Staphylococcus_MecI                YKKGFIDRKK----DNKIF--QYYSLVEESDIKYKTSKNFINKVYKGGFNSLVLNFVEK-
Staphylococcus_BlaI                YKKEIIKRYK----SENIY--FYSSNIKEDDIKMKTAKTFLNKLYGGDMKSLVLNFAKN-
                                    .*  :   .      .     : . :.  :     :   :: .      . *  * .

Leptolyngbya_sp_PCC_7375           DQDSLDQLEAITKRIQAARRARSNNQQGQEK
Pseud0anabaena_sp_PCC_6802         DLASVDRLEAIAQRLKAIRQEREER------
Synechocystis                      DTASIDQLTAIADRLRAARQQRQEEK-----
Synechococcus_sp_JA-2-3B'a(2-13)   DQASVSQLQAIAERLRAARQARRDPR-----
Acaryochloris_marina_MBIC11017     DQASVEQLEEIADKIRAARKAREDQ------
Anabaena7120                       DEAASEQIEAIAKRIQAARQAREEQ------
Bacillus_licheniformis_BlaI        DQLSGEEINELYQILEEHKNRKKE-------
Staphylococcus_MecI                EDLSQDEIEELRNILNKK-------------
Staphylococcus_BlaI                EELNNKEIEELRDILNDISKK----------
                                   :    . :  : . :
```

**Figure S7. Alignment of cyanobacterial PetR proteins with *Staphylococcus* and *Bacillus* BlaI/MecI proteins.** Protein sequences were aligned using MUSCLE using the default parameters Protein sequences were aligned using MUSCLE using the default parameters. The initial degradation site is highlighted in red.

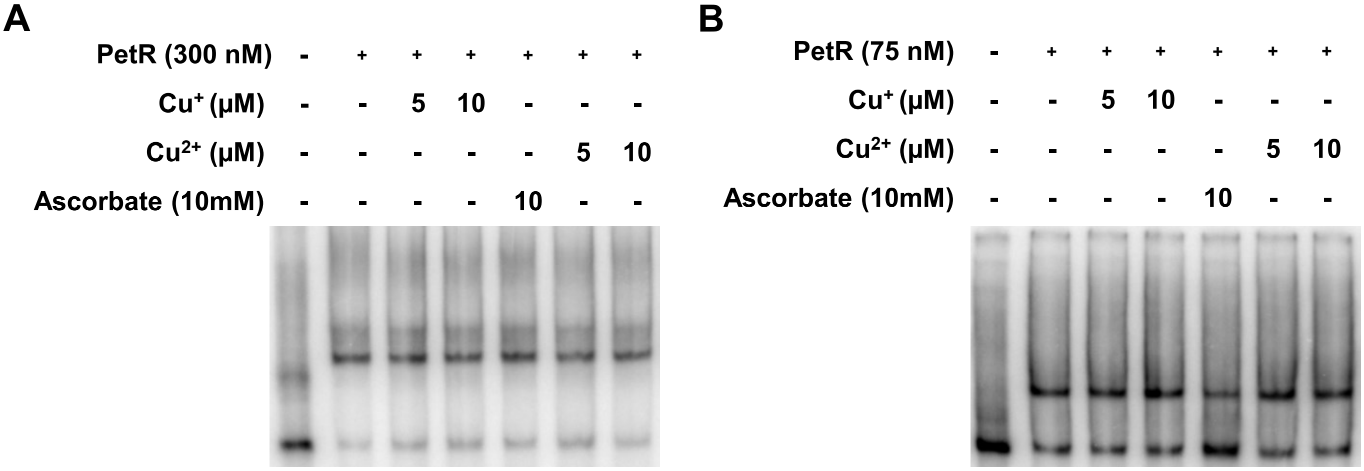


**Figure S8. PetR binding to *petE* and *petJ* promoters is not affected by copper**.

A. Binding of recombinant PetR (300 nM) to a 103 bp *petJ* promoter probe in the presence of 5 or 10 μM of $Cu^{2+}$ or $Cu^{+}$ (reduced by 1000-fold excess of ascorbate). A sample containing only ascorbate was included as a control.

B. Binding of recombinant PetR (50 nM) to a 217 bp *petE* promoter probe in the presence of 5 or 10 μM of $Cu^{2+}$ or $Cu^{+}$ (reduced by 1000 fold excess of ascorbate). A sample containing only ascorbate (10 mM) was included as a control.

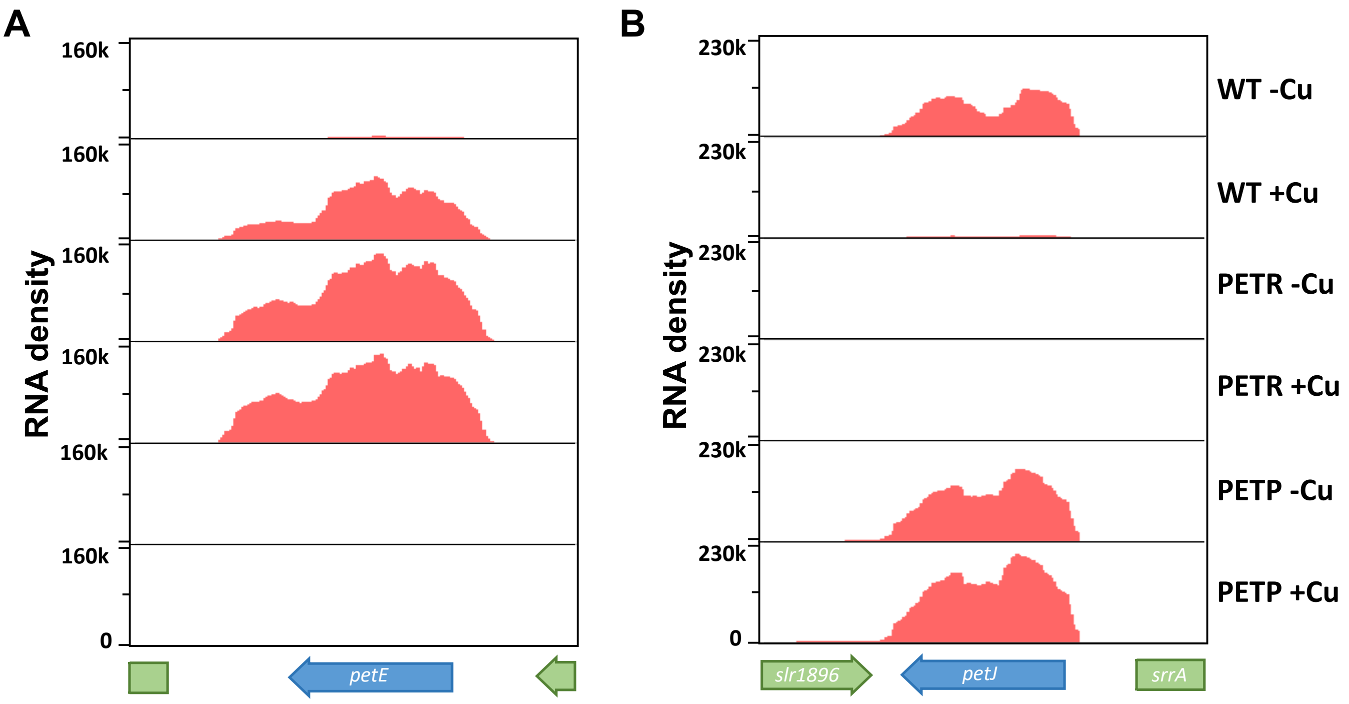


**Figure S9. RNA-seq analysis of WT, PETR and PETP after copper addition**.

A. RNA seq density profile of *petE* in WT, PETR and PETP before and after 2h of 0.5 μM copper addition.
B. RNA seq density profile of *petJ* in WT, PETR and PETP before and after 2h of 0.5 μM copper addition.

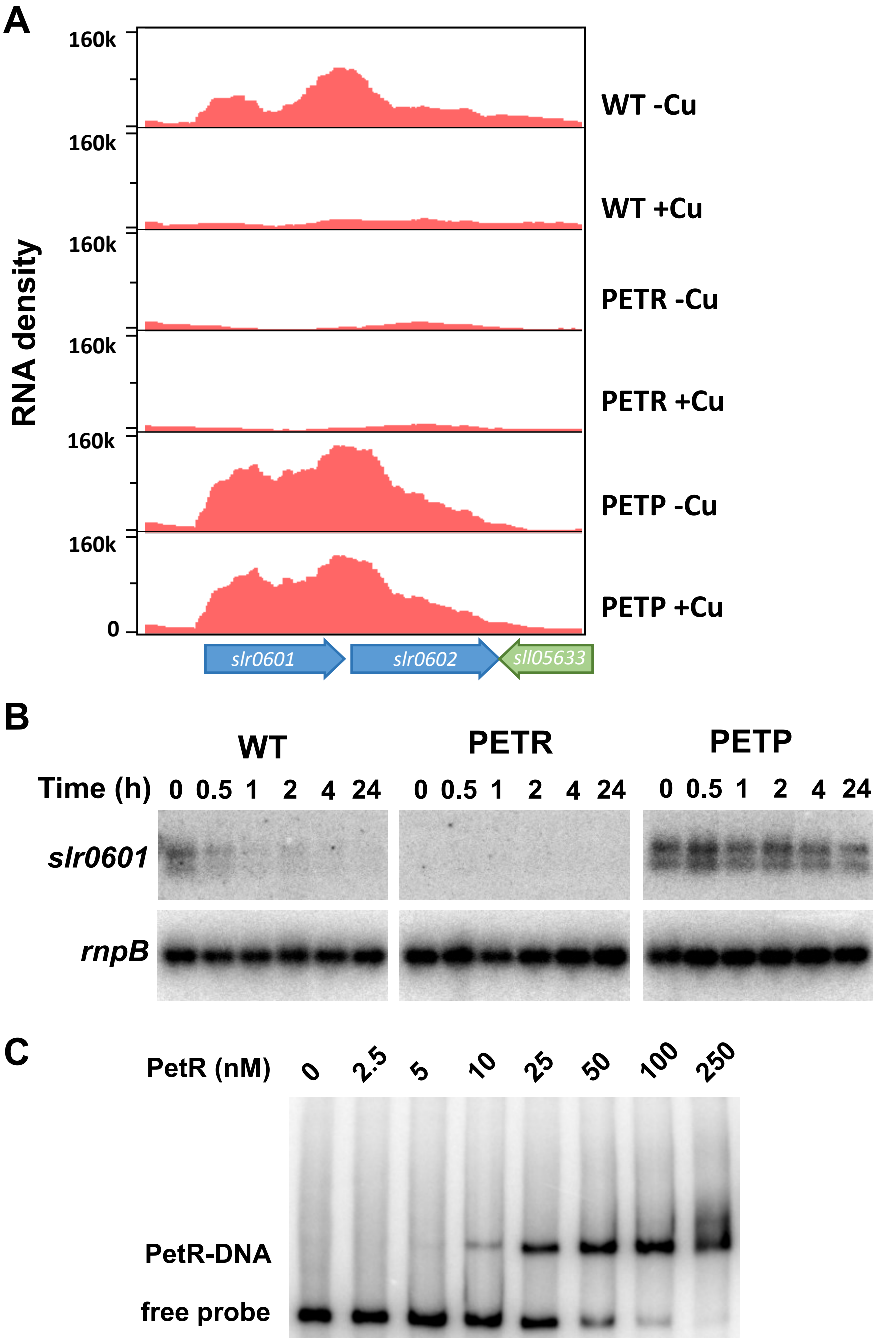


**D**

**cccgcaaatggattcattagataaattattggcacataaaattagtggactttcgccct**
**agcccgccgggaaaccaagtttgacgggctgtctttttgacccttgaacgttattctgcc**
**taggataggggGcattcttttagcagaaatttactaaatttccccATG**

**Figure S10. PetR binds to *slr0601* promoter.**

A. RNA seq density profile of *slr0601* and *slr0602* in WT, PETR and PETP strains before and after 2h of 0.5 μM copper addition.
B. RNA blot of *slr0601* in WT, PETR and PETP strains in response to 0.5 μM copper addition. Total RNA was isolated from cells grown in BG11C-Cu medium at the indicated times after addition of 0.5 μM of copper .The filters were hybridized with *slr0601* probe and subsequently stripped and re-hybridized with a *rnpB* probe as a control.
C. Binding of recombinant PetR to a 177 bp *slr0601* promoter probe. The indicated PetR concentration were used in each lane.
*D.* *slr0601* promoter sequence. Transcriptional start site is in capital letters and underlined, -10 boxes are underlined and conserved nucleotides from the motif found in Figure 4c is in purple. Direct repeats identified in Figure 3D are in red.

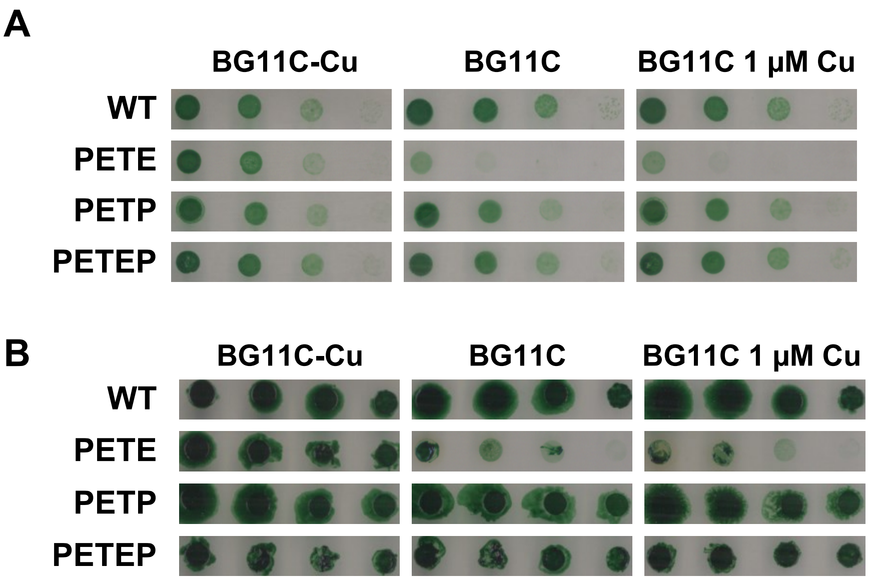


**Figure S11. PETE strain is able to grow at reduced rates in the presence of copper.**

A. Growth of WT, PETE, PETP, PETEP strains in different copper availability regimes. Tenfold serial dilutions of a 1 µg chlorophyll $mL^{-1}$ cells suspension were spotted onto BG11C-Cu, BG11C-Cu o BG11C+ 1 µM Cu. Plates were photographed after 5 days of growth.

B. Growth of WT, PETE, PETP, PETEP strains in different copper availability regimes. Tenfold serial dilutions of a 1 µg chlorophyll $mL^{-1}$ cells suspension were spotted onto BG11C-Cu, BG11C-Cu o BG11C+ 1 µM Cu. Plates were photographed after 10 days of growth.

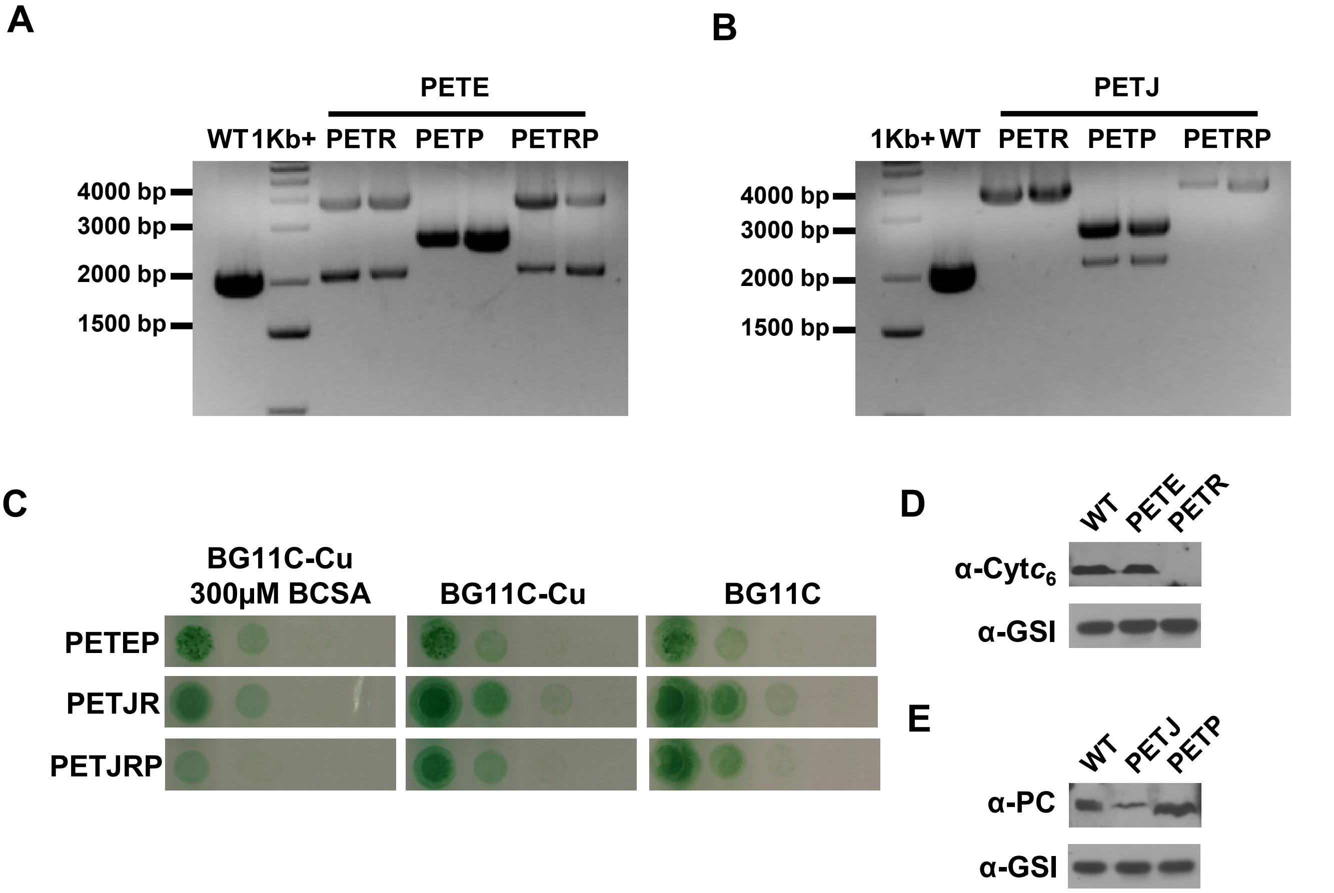


**Figure S12. Genetic interactions between *petJ*, *petE* and *petRP*.**

A. PCR analysis of the *petRP* locus in WT, PETER, PETEP and PETERP strains (two clones were analysed for each strain) using oligonucleotides 223-249.

B. PCR analysis of the *petRP* in WT, PETJR, PETJP and PETJRP mutant strains (two clones were analysed for each strain) using oligonucleotides 223-249.

C. Growth of PETEP, PETJR and PETJRP strains in different copper availability regimes. Tenfold serial dilutions of a 1 µg chlorophyll $mL^{-1}$ cells suspension were spotted onto BG11C-Cu+BCSA, BG11C-Cu o BG11C. Plates were photographed after 5 days of growth.

D. Western blot analysis of $Cytc_6$ levels in WT, PETE and PETR strains. Cells were grown in BG11C medium and harvested during exponential growth (3-5 µg chl $mL^{-1}$). 150 µg of total protein from soluble extracts was separated by 15 % SDS-PAGE and subjected to western blot to detect Cyt$c_6$ or GSI.

E. Western blot analysis of PC levels in WT, PETJ and PETP strains. Cells were grown in BG11C-Cu+BCSA medium and harvested during exponential growth (3-5 µg chl $mL^{-1}$). 150 µg of total protein from soluble extracts was separated by 15 % SDS-PAGE and subjected to western blot to detect PC or GSI.

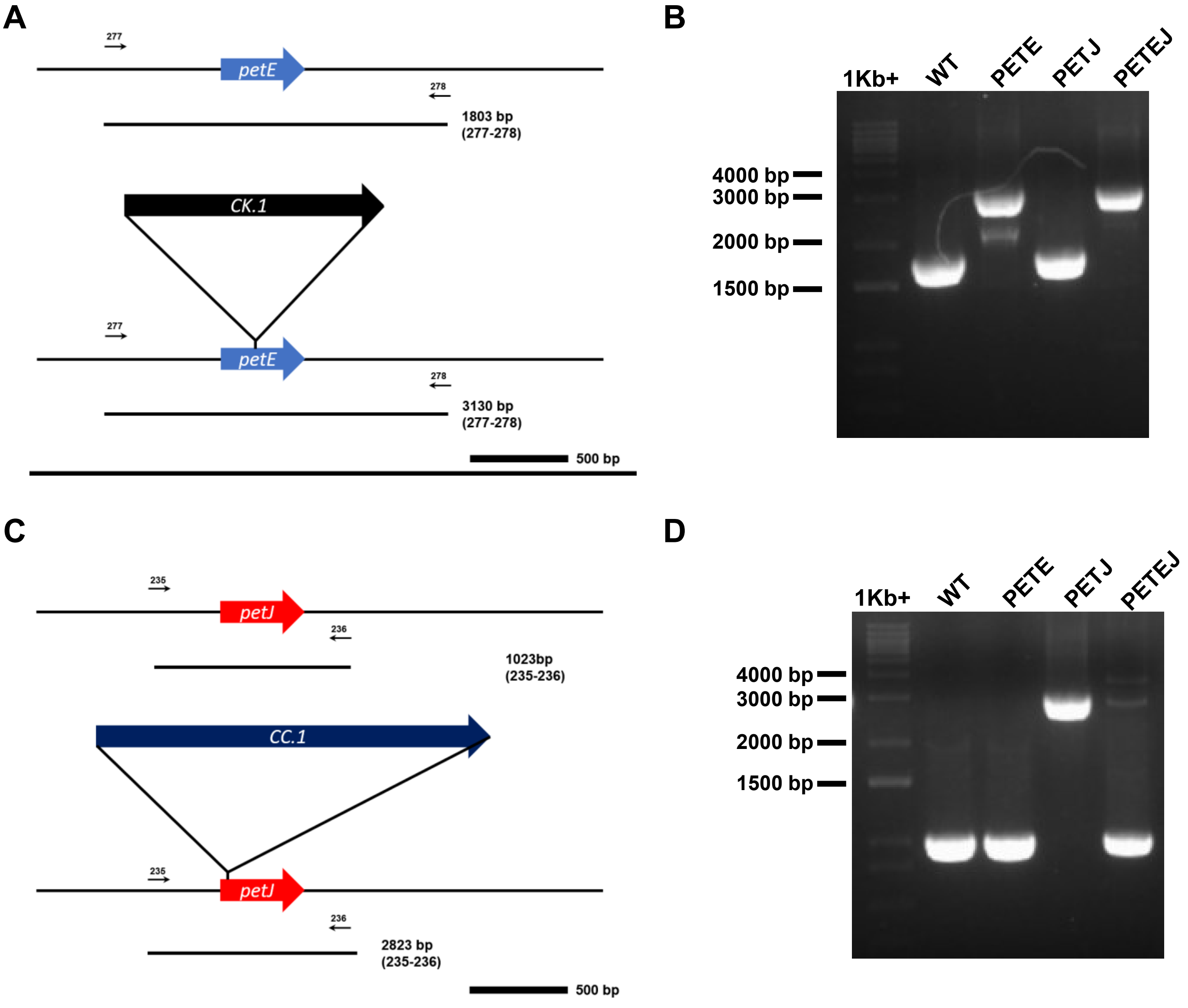


**Figure S13. Construction of PETEJ double mutants.**

A. Schematic representation of the *petE* genomic region in WT and PETE strains. Oligonucleotides and sizes of bands amplified by the different oligo pairs in all strains are shown.
B. PCR analysis of the *petE* locus in WT, PETE, PETJ and PETEJ strains using oligonucleotides 174-175.
C. Schematic representation of the *petJ* genomic region in WT and PETJ strains. Oligonucleotides and sizes of bands amplified by the different oligo pairs in all strains are shown
D. PCR analysis of the *petJ* locus in WT, PETE, PETJ and PETEJ strains using oligonucleotides 235-236.

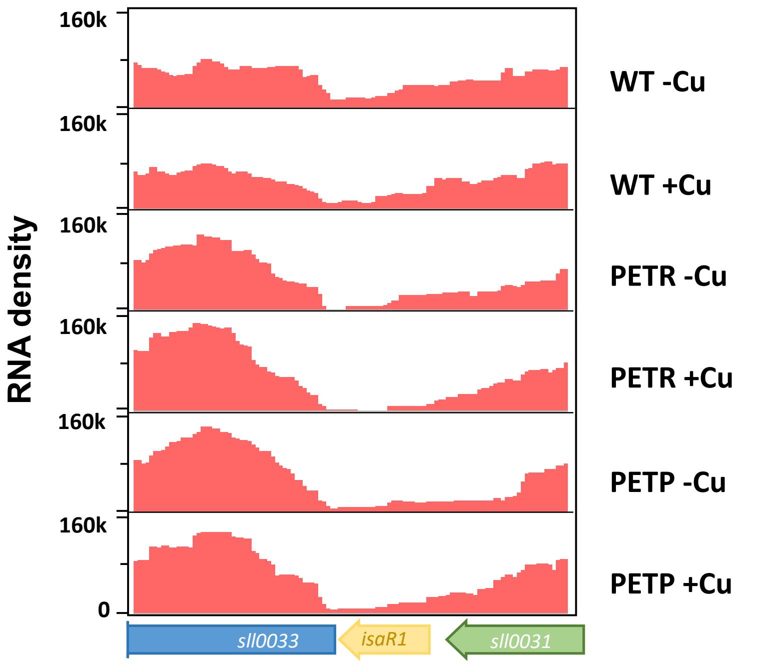


**Figure S14. *isaR* ncRNA is not regulated by *petRP*.**
RNA seq density profile of *isaR* in WT, PETR and PETP strains before and after 2h of 0.5 µM copper addition.

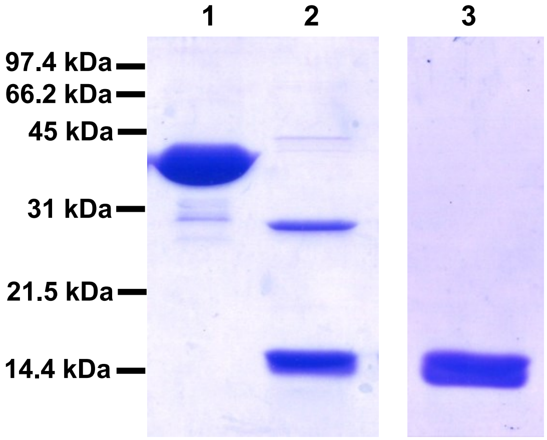


**Figure S15. SDS-Page of recombinant PetR versions.**
5 µg of purified GST-PetR (1), GST-PetR after in column digestion with Precission protease (2) or PetR after Heparin purification (3) were separated in 12% SDS-PAGE and gels were stained with Coomassie Brilliant Blue R-250.

**Table S1. Differentially regulated genes ($P_{adj}$ <0.01, 2- fold change (n= 3)) identified in WT, PETR and PETP strains after copper addition. NS: not significant; repressed genes are expressed as negative fold change.**

| systematic ID | Fold Change WT | Fold Change PETR | Fold Change PETP | Gene name | Annotation |
|---|---|---|---|---|---|
| sll0199 | 24.28 | NS | NS | *petE* | plastocyanin |
| sll0788 | 9.40 | 10.88 | 9.06 | *copM* | copper binding protein involves in copper resistance; chromosomal copy |
| sll0789 | 16.58 | 18.35 | 31.88 | *copR, rre34* | two-component response regulator OmpR subfamily involved in copper resistance; chromosomal copy |
| sll0790 | 6.16 | 5.00 | 8.08 | *hik31, copS* | two-component sensor histidine kinase involved in copper resistance; chromosomal copy |
| sll1796 | -16.66 | NS | NS | *petJ* | cytochrome $c_6$ |
| slr0601 | -5.55 | NS | NS | *slr0601* | unknown protein |
| slr0602 | -2.32 | NS | NS | *slr0602* | unknown protein |
| slr0793 | 6.50 | 13.51 | 9.34 | *nrsB* | cation efflux system protein involved in nickel and cobalt tolerance |
| slr0794 | 4.29 | 5.66 | 5.01 | *nrsA* | cation efflux system protein involved in nickel and cobalt tolerance |
| slr0795 | 5.10 | 5.46 | 7.46 | *nrsC* | cation efflux system protein involved in nickel and cobalt tolerance |
| slr6039 | 11.75 | 14.26 | 11.98 | *pcopM,hypoP* | copper binding protein involves in copper resistance; plasmid copy |

| systematic ID | Fold Change WT | Fold Change PETR | Fold Change PETP | Gene name | Annotation |
|---|---|---|---|---|---|
| slr6040 | 16.16 | 16.19 | 40.61 | *pcopR, rreP* | two-component response regulator OmpR subfamily involved in copper resistance; plasmid copy |
| slr6041 | 10.20 | 7.53 | 9.07 | *pcopS, hikP,* | two-component sensor histidine kinase involved in copper resistance; plasmid copy |
| slr6042 | 18.47 | 21.36 | 27.93 | *copB* | cation efflux system protein involved in copper tolerance; czcB homolog |
| slr6043 | 17.30 | 23.26 | 29.62 | *copA* | cation efflux system protein involved in copper tolerance; czcA homolog |
| slr6044 | 9.56 | 14.28 | 16.85 | *copC* | cation efflux system protein involved in copper tolerance; |
| slr6045 | 4.37 | 6.13 | 6.20 | *slr6045* | unknown protein |
| slr0145 | NS | -2.44 | NS | *slr0145* | unknown protein |
| slr0917 | NS | NS | -2.0 | *bioF* | 7-keto-8-aminopelargonic acid synthetase |

Table S2. *Synechocystis* strains used in this work.

| STRAIN | RELEVANT GENOTYPE | Mutated ORFs | SOURCE |
|---|---|---|---|
| **WT** | *Synechocystis* sp. PCC 6803 | - | Lab collection |
| **PETE** | *petE::C.K1* | *sll0199* | *(Giner-Lamia et al., 2012)* |
| **PETJ** | *petJ::Cm* | *sll1796* | This study |
| **PETR** | *petR::Sp* | *slr0240* | This study |
| **PETR2** | *petR::Sp glnN::*$P_{petR}$*RP:Nat* | *slr0240*, *slr0288* | This study |
| **PETR·** | *petR::Sp glnN::*$P_{cpcB}$*RP:Nat* | *slr0240*, *slr0288* | This study |
| **PETP** | Δ*petP::Ery* | *slr0241* | This study |
| **PETRP** | Δ*petPR::Sp* | *slr0240*, *slr0241* | This study |
| **PETER** | *petE::C.K1; petR::Sp* | *sll0199*, *slr0240* | This study |
| **PETJR** | *petJ::Cm; petR::Sp* | *sll1796*, *slr0240* | This study |
| **PETEP** | *petE::C.K1;* Δ*petP::Ery* | *sll0199*, *slr0241* | This study |
| **PETJP** | *petJ::Cm;* Δ*petP::Ery* | *sll1796*, *slr0241* | This study |
| **PETERP** | *petE::C.K1;* Δ*petRP::Sp* | *sll0199*, *slr0240*, *slr0241* | This study |
| **PETJRP** | *petJ::Cm;* Δ*petRP::Sp* | *sll1796*, *slr0240*, *slr0241* | This study |

| STRAIN | RELEVANT GENOTYPE | Mutated ORFs | SOURCE |
|---|---|---|---|
| **PETER** | *petE::C.K1; petR::Sp* | *sll0199, slr0240* | This study |
| **PETEJ** | *petE::C.K1; petJ::Cm* | *sll0199, sll1796* | This study |

**Table S3. Oligonucleotides used in this work.**

| Number | Name | Sequence | Used for |
|---|---|---|---|
| 41 | Prom_Ck1_R | AAGAGTGGTACCCATGGTAA ACGATCCTCATCCTGTCTC | PETR2 and PETR3 segregation |
| 50 | glnN_check_F | ATGCAGGCCAGTCTTCCTAA | PETR2 and PETR3 segregation |
| 51 | glnN_check_R | AAATGGCAGTGTCCAAGTCC | PETR2 and PETR3 segregation |
| 53 | Prom_petE_N otI | cggcggccgcTGAGGCTGTATAA TCTACG | Band-shift assay probe |
| 92 | petE_ATG_Sal | CAGTCGACTTGGCGATTGTA TCTATAGGG | Band-shift assay probe |
| 223 | copY_KO_F | CCAAGGCAAGATTGTGGTGC | For pPETR plasmid construction |
| 224 | copY_KO_R | CCTGTTCCGGAGACAGCAAT | For pPETR plasmid construction |
| 248 | slr0241_check _F | CAACAGCGACAGGAGGAGAA | For pPETP plasmid construction |
| 249 | slr0241_check _R | AAAGCTCCGGTTGAGAAGGG | For pPETP plasmid construction |
| 235 | petJ KO F | CGAGACGGCACTGAGGATTT | For pPETJ plasmid construction |
| 236 | petJ KOR | AGTTGGGCTTTACGGGCTAC | For pPETP plasmid construction |
| 256 | copY_comple mentation_Fw | gactgaattcCCCCGGCGATCGC GCCTTTATC | For pPETRComp and pNPETRP plasmid construction |
| 257 | slr0241_R_Xh oI | atcgctcgag TCACCTGAATAGCTTGAAC | For pNPETRP plasmid construction |
| 258 | copY_OE_Fw_ SalI | gtcgacGGAGGTtatcattATGAGC TGGATTCCCCCC | For pPETR_OE plasmid construction |

| Number | Name | Sequence | Used for |
|---|---|---|---|
| 233 | copY F NcoI BHI | ccatgg ggatccATGAGCTGGATTCCCC CCTAC | pGEX_PETR plasmid construction |
| 234 | copY R NotI | atgcggccgcAATACTGTGCATTA TTTCTCC | pGEX_PETR, pPETR_comp and pPETR_OE plasmid construction |
| 172 | petEF | acaatcctcgctggccttct | *petE* probe |
| 173 | petER | cgacaactttgcctaccatg | *petE* probe |
| 174 | petJF | attcaaccaagctagccgaa | *petJ* probe |
| 175 | petJR | tccgcttgatcaagcacgta | *petJ* probe |
| 178 | rnpB_LEFT | GAGTTGCGGATTCCTGTCAC | *rnpB* probe |
| 179 | rnpB_RIGHT | AATTCCTCAAGCGGTTCCAC | *rnpB* probe |
| 275 | slr0601F | gtctttagactgcagcagcc | *slr0601* probe |
| 276 | slr0601R | tgattgctgtgctccatggg | *slr0601* probe |
| 389 | COPM1F | agcattcccatgggtaatcaattctggatat | *copM* probe |
| 390 | COPM1R | catctcgagtcactgaccataccagttttga ta | *copM* probe |
| 299 | oligo F1_new_petJS alI | GATCGTCGACccacttcacaaacct agccaat | Band-shift assay probe |
| 262 | oligo R3 | cgcgaaggtattatgggaggcggtccagg caaa | Band-shift assay probe |
| 295 | slr0601_retard o_F_SalI | gccaGTCGACcccgcaaatggattcat tag | Band-shift assay probe |
| 296 | slr0601_retard o_R_NotI | acatGCGGCCGCttagtaaatttctgct aaaag | Band-shift assay probe |

Dataset S1. List of cyanobacteria strains used for phylogenomic analysis obtained from NCBI and accession numbers for *petJ*, *petE* and *petRP* genes.
Dataset S2. Promoter sequences used for MEME analysis
Dataset S3. Calculated RPKM for all *Synechocystis* genes.
Dataset S4. Differentially expressed genes identified using RNA-seq.
