## Supplementary Figures for "A protease-mediated mechanism regulates the cytochrome *c*_6_/ plastocyanin switch in *Synechocystis* sp. PCC 6803"

### Slide 1
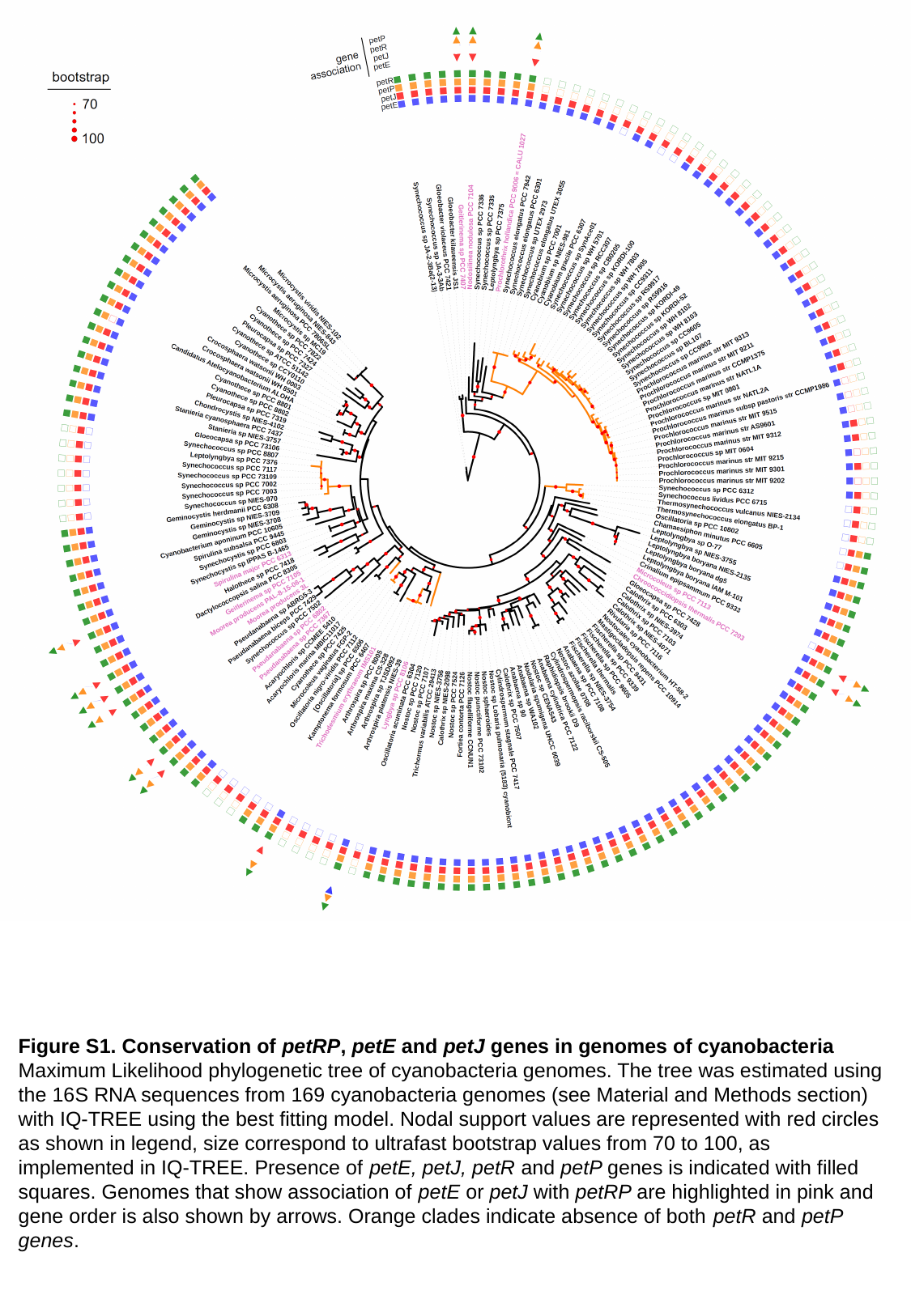

Figure S1. Conservation of petRP, petE and petJ genes in genomes of cyanobacteria
Maximum Likelihood phylogenetic tree of cyanobacteria genomes. The tree was estimated using the 16S RNA sequences from 169 cyanobacteria genomes (see Material and Methods section) with IQ-TREE using the best fitting model. Nodal support values are represented with red circles as shown in legend, size correspond to ultrafast bootstrap values from 70 to 100, as implemented in IQ-TREE. Presence of petE, petJ, petR and petP genes is indicated with filled squares. Genomes that show association of petE or petJ with petRP are highlighted in pink and gene order is also shown by arrows. Orange clades indicate absence of both petR and petP genes.

### Slide 2
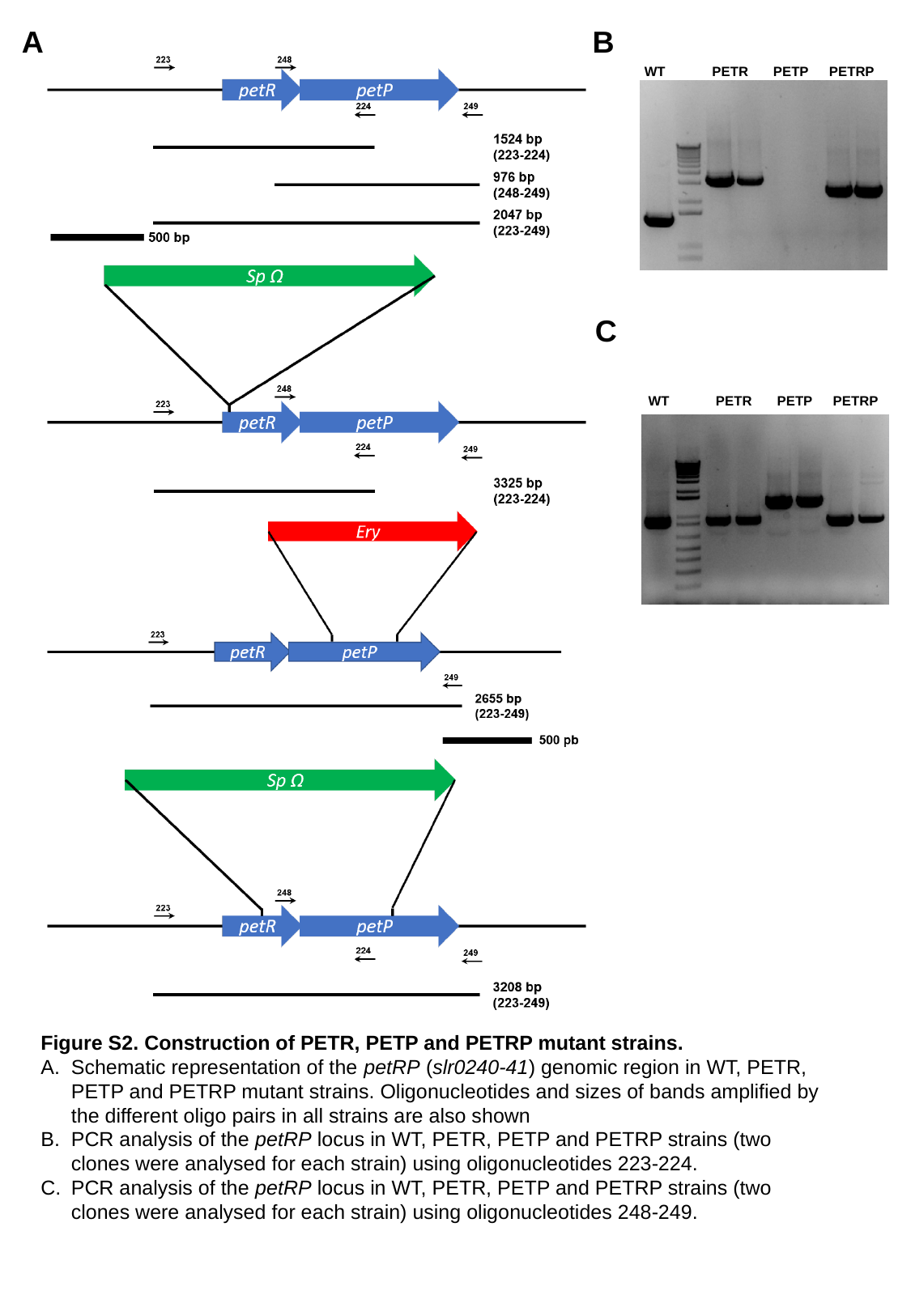

A
B
WT
PETR
PETP
PETRP
C
WT
PETR
PETP
PETRP
Figure S2. Construction of PETR, PETP and PETRP mutant strains.
Schematic representation of the petRP (slr0240-41) genomic region in WT, PETR, PETP and PETRP mutant strains. Oligonucleotides and sizes of bands amplified by the different oligo pairs in all strains are also shown
PCR analysis of the petRP locus in WT, PETR, PETP and PETRP strains (two clones were analysed for each strain) using oligonucleotides 223-224.
PCR analysis of the petRP locus in WT, PETR, PETP and PETRP strains (two clones were analysed for each strain) using oligonucleotides 248-249.

### Slide 3
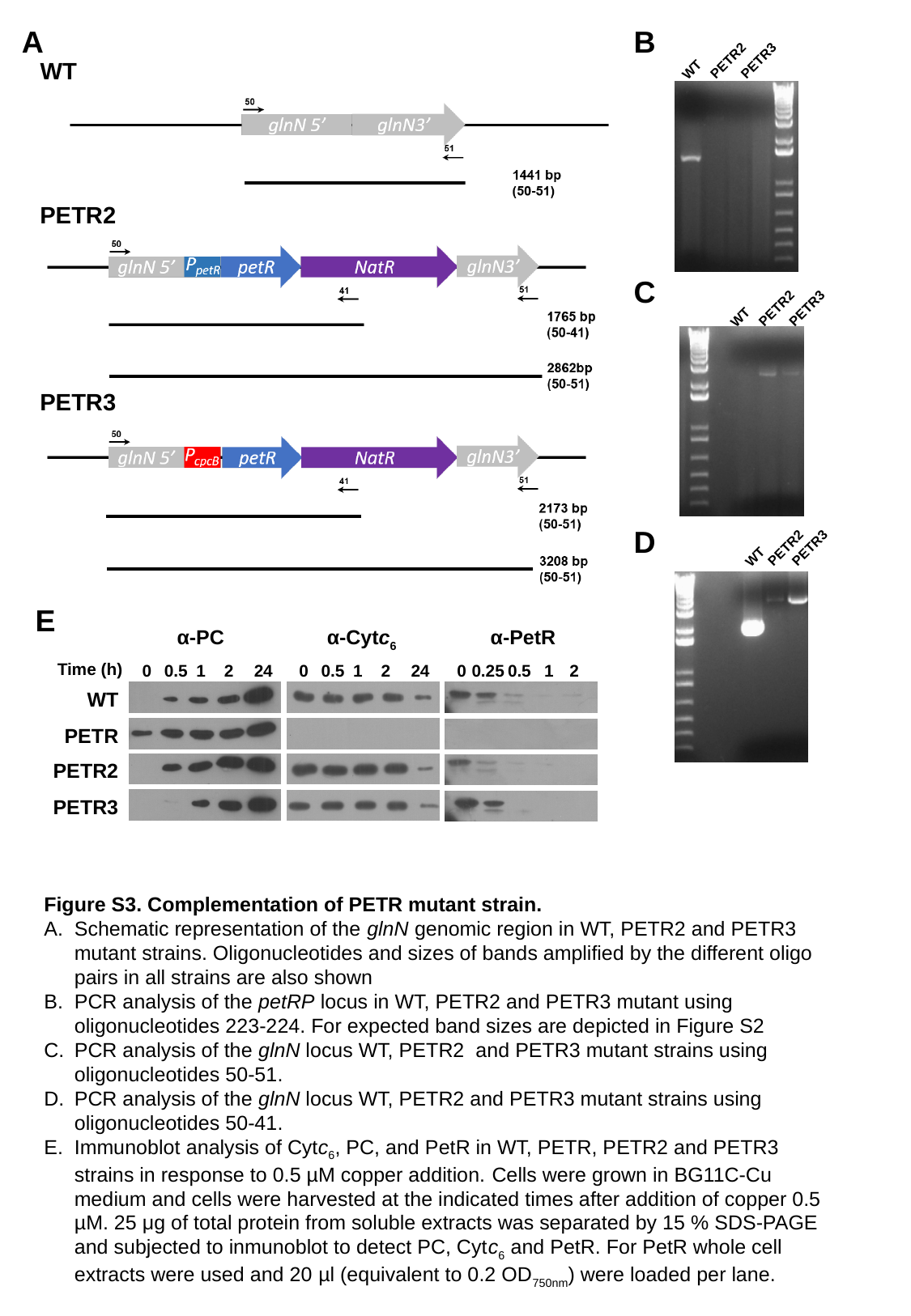

A
B
PETR2
PETR3
WT
WT
PETR2
C
PETR2
PETR3
WT
PETR3
D
PETR2
PETR3
WT
E
α-PC
α-Cytc_6_
α-PetR
Time (h)
0
0.5
1
2
24
0
0.5
1
2
24
0
0.25
0.5
1
2
WT
PETR
PETR2
PETR3
Figure S3. Complementation of PETR mutant strain.
Schematic representation of the glnN genomic region in WT, PETR2 and PETR3 mutant strains. Oligonucleotides and sizes of bands amplified by the different oligo pairs in all strains are also shown
PCR analysis of the petRP locus in WT, PETR2 and PETR3 mutant using oligonucleotides 223-224. For expected band sizes are depicted in Figure S2
PCR analysis of the glnN locus WT, PETR2 and PETR3 mutant strains using oligonucleotides 50-51.
PCR analysis of the glnN locus WT, PETR2 and PETR3 mutant strains using oligonucleotides 50-41.
Immunoblot analysis of Cytc_6_, PC, and PetR in WT, PETR, PETR2 and PETR3 strains in response to 0.5 µM copper addition. Cells were grown in BG11C-Cu medium and cells were harvested at the indicated times after addition of copper 0.5 µM. 25 μg of total protein from soluble extracts was separated by 15 % SDS-PAGE and subjected to inmunoblot to detect PC, Cytc_6_ and PetR. For PetR whole cell extracts were used and 20 µl (equivalent to 0.2 OD_750nm_) were loaded per lane.

### Slide 4
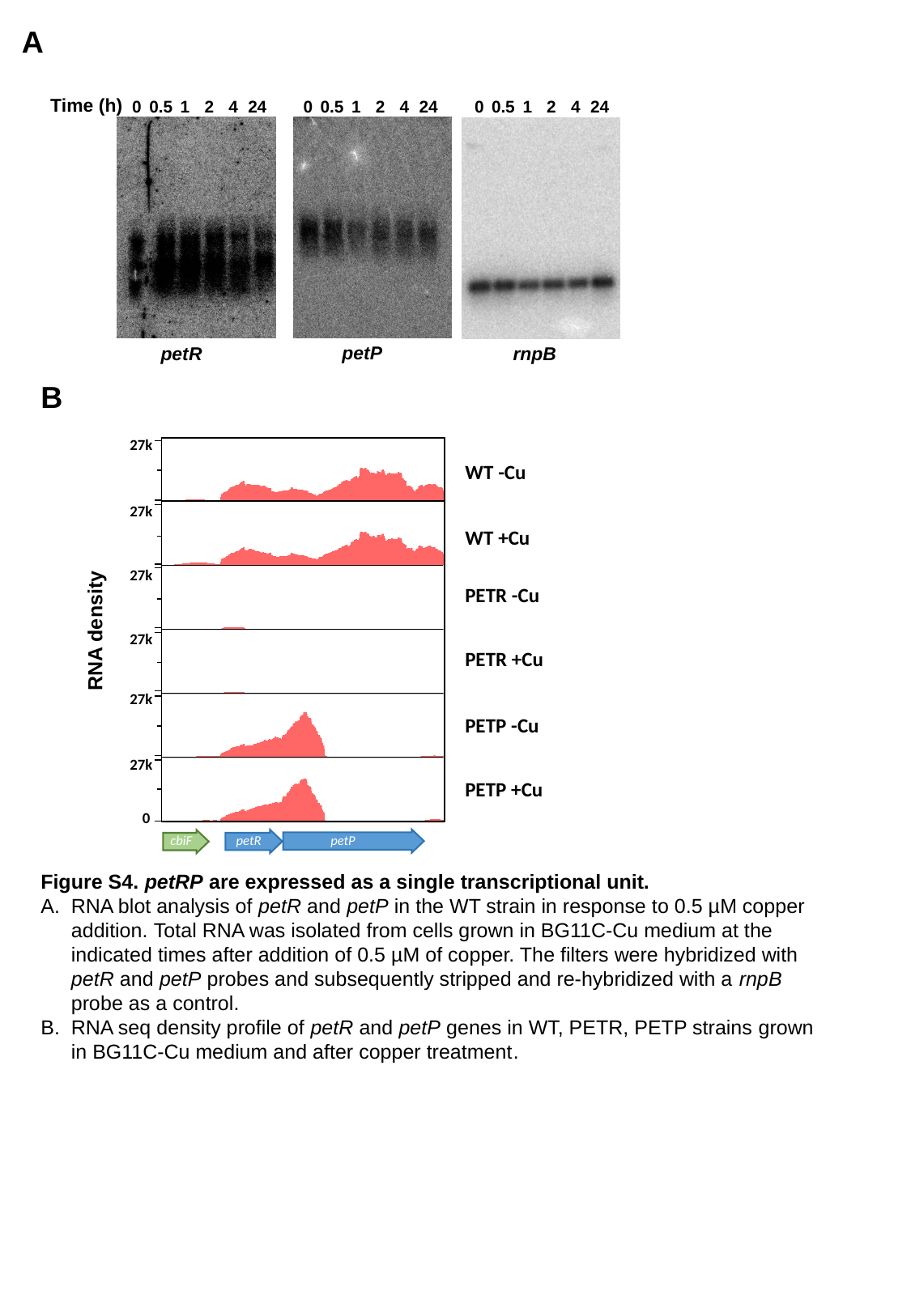

A
Time (h)
0
0.5
1
2
4
24
0
0.5
1
2
4
24
0
0.5
1
2
4
24
petP
petR
rnpB
B
27k
WT -Cu
27k
WT +Cu
27k
PETR -Cu
RNA density
27k
PETR +Cu
27k
PETP -Cu
27k
PETP +Cu
0
cbiF
petR
petP
Figure S4. petRP are expressed as a single transcriptional unit.
RNA blot analysis of petR and petP in the WT strain in response to 0.5 µM copper addition. Total RNA was isolated from cells grown in BG11C-Cu medium at the indicated times after addition of 0.5 µM of copper. The filters were hybridized with petR and petP probes and subsequently stripped and re-hybridized with a rnpB probe as a control.
RNA seq density profile of petR and petP genes in WT, PETR, PETP strains grown in BG11C-Cu medium and after copper treatment.

### Slide 5
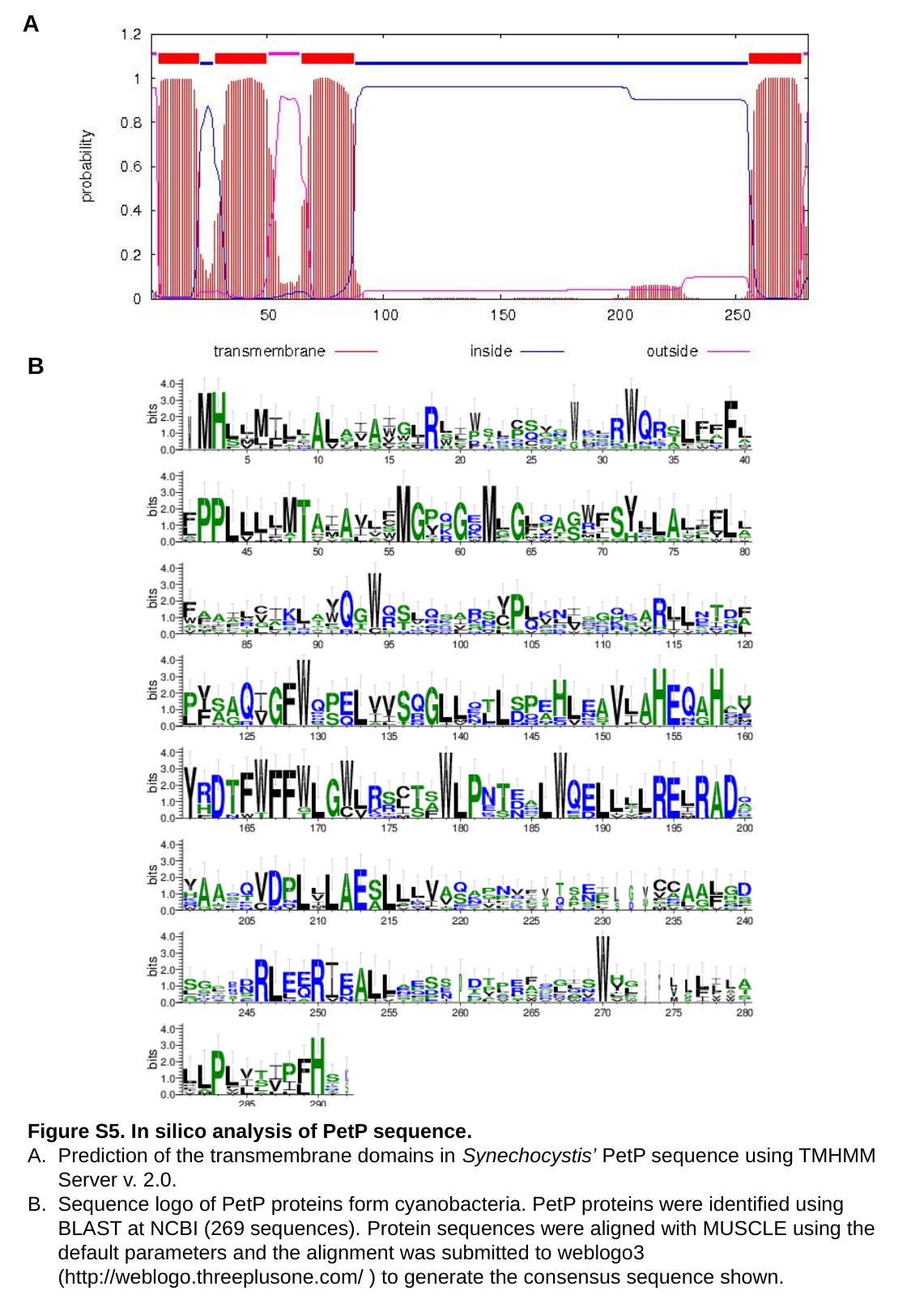

A
B
Figure S5. In silico analysis of PetP sequence.
Prediction of the transmembrane domains in Synechocystis’ PetP sequence using TMHMM Server v. 2.0.
Sequence logo of PetP proteins form cyanobacteria. PetP proteins were identified using BLAST at NCBI (269 sequences). Protein sequences were aligned with MUSCLE using the default parameters and the alignment was submitted to weblogo3 (http://weblogo.threeplusone.com/ ) to generate the consensus sequence shown.

### Slide 6
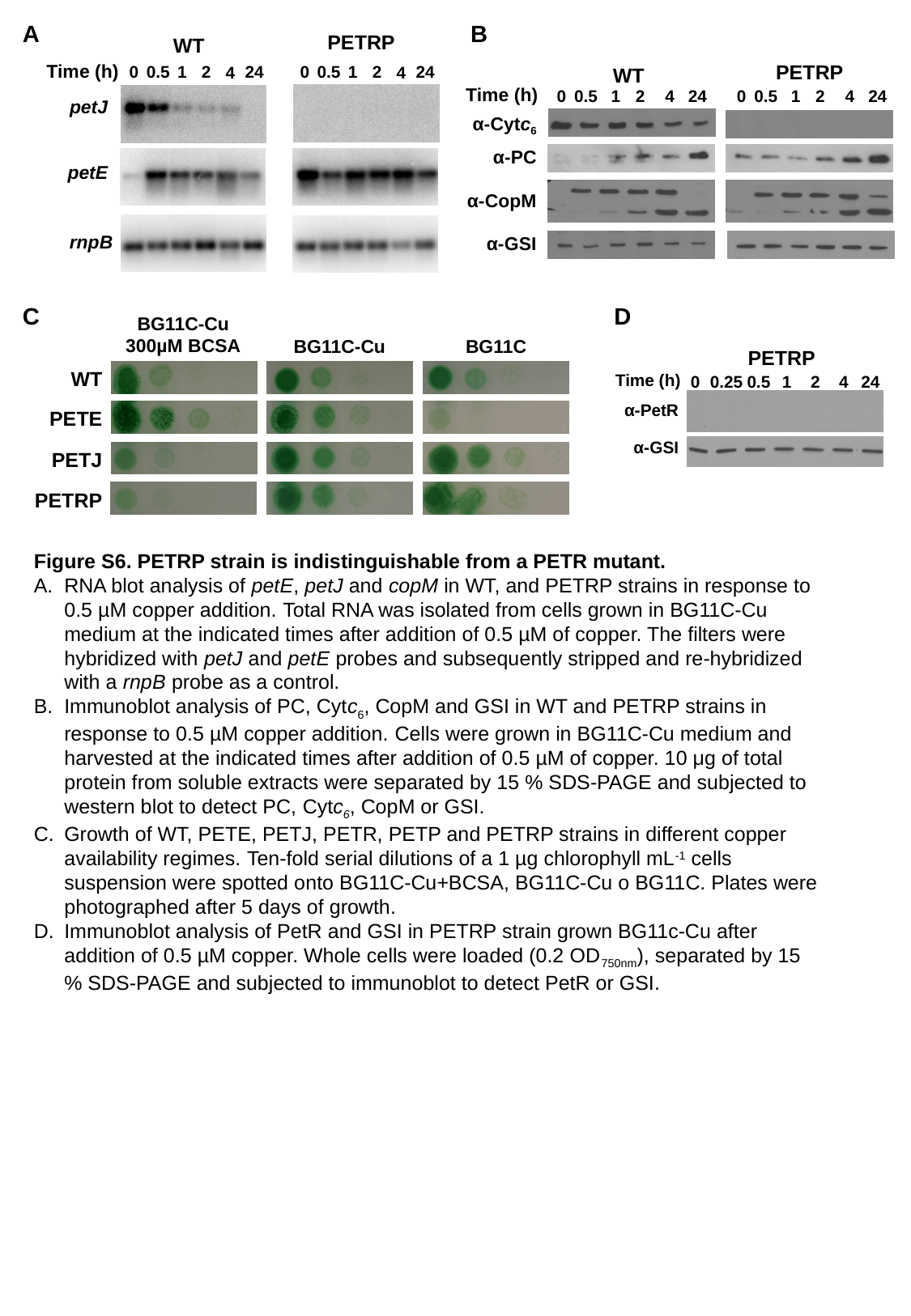

A
B
PETRP
WT
Time (h)
0
0.5
1
2
24
4
0
0.5
1
2
24
4
petJ
petE
rnpB
PETRP
WT
Time (h)
0
0.5
1
2
4
24
0
0.5
1
2
4
24
α-Cytc_6_
α-PC
α-CopM
α-GSI
C
D
BG11C-Cu
300µM BCSA
BG11C-Cu
BG11C
WT
PETE
PETJ
PETRP
PETRP
Time (h)
0
0.25
0.5
1
2
4
24
α-PetR
α-GSI
Figure S6. PETRP strain is indistinguishable from a PETR mutant.
RNA blot analysis of petE, petJ and copM in WT, and PETRP strains in response to 0.5 µM copper addition. Total RNA was isolated from cells grown in BG11C-Cu medium at the indicated times after addition of 0.5 µM of copper. The filters were hybridized with petJ and petE probes and subsequently stripped and re-hybridized with a rnpB probe as a control.
Immunoblot analysis of PC, Cytc_6_, CopM and GSI in WT and PETRP strains in response to 0.5 µM copper addition. Cells were grown in BG11C-Cu medium and harvested at the indicated times after addition of 0.5 µM of copper. 10 μg of total protein from soluble extracts were separated by 15 % SDS-PAGE and subjected to western blot to detect PC, Cytc_6_, CopM or GSI.
Growth of WT, PETE, PETJ, PETR, PETP and PETRP strains in different copper availability regimes. Ten-fold serial dilutions of a 1 µg chlorophyll mL^-1^ cells suspension were spotted onto BG11C-Cu+BCSA, BG11C-Cu o BG11C. Plates were photographed after 5 days of growth.
Immunoblot analysis of PetR and GSI in PETRP strain grown BG11c-Cu after addition of 0.5 µM copper. Whole cells were loaded (0.2 OD_750nm_), separated by 15 % SDS-PAGE and subjected to immunoblot to detect PetR or GSI.

### Slide 7
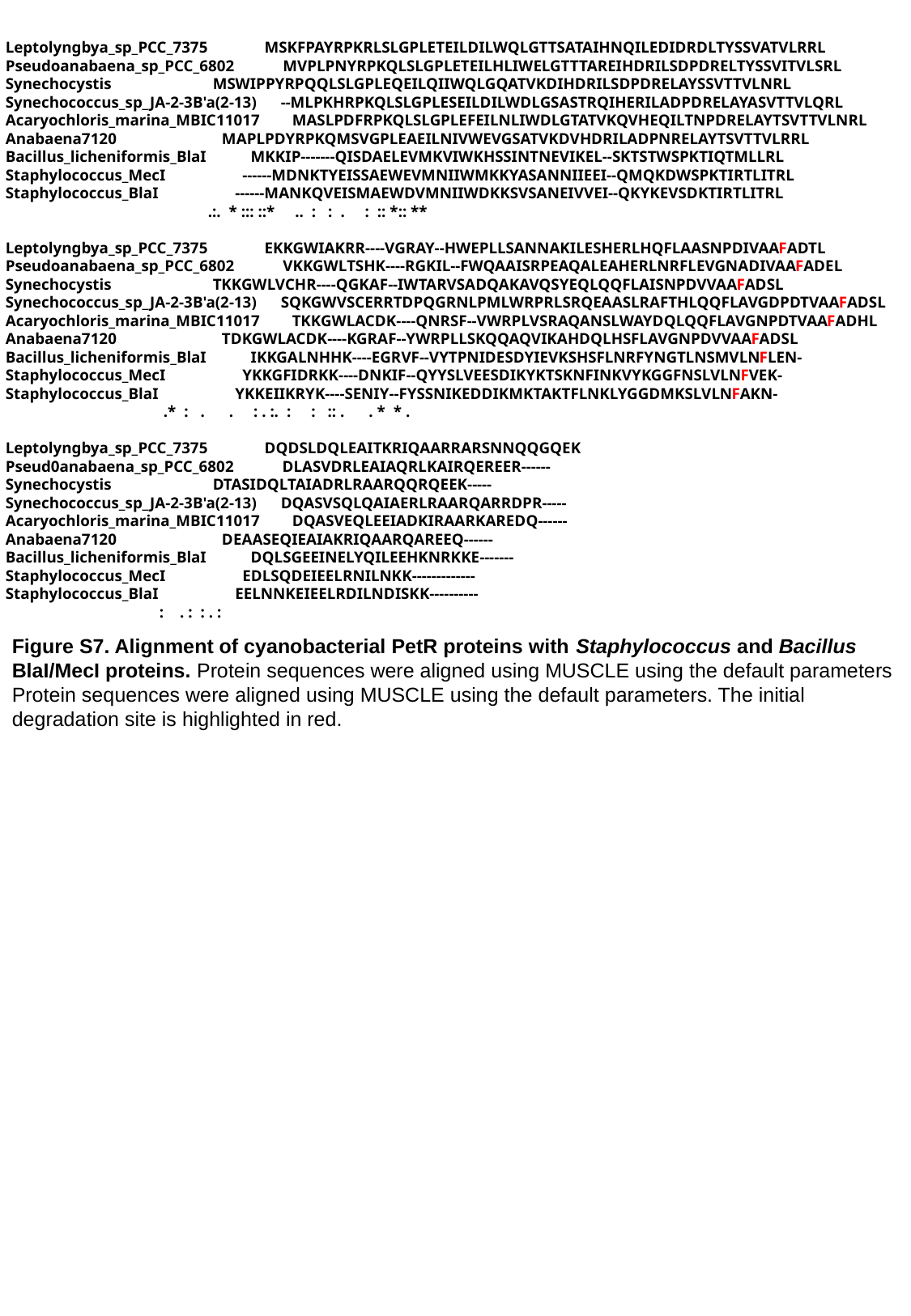

Leptolyngbya_sp_PCC_7375 MSKFPAYRPKRLSLGPLETEILDILWQLGTTSATAIHNQILEDIDRDLTYSSVATVLRRL
Pseudoanabaena_sp_PCC_6802 MVPLPNYRPKQLSLGPLETEILHLIWELGTTTAREIHDRILSDPDRELTYSSVITVLSRL
Synechocystis MSWIPPYRPQQLSLGPLEQEILQIIWQLGQATVKDIHDRILSDPDRELAYSSVTTVLNRL
Synechococcus_sp_JA-2-3B'a(2-13) --MLPKHRPKQLSLGPLESEILDILWDLGSASTRQIHERILADPDRELAYASVTTVLQRL
Acaryochloris_marina_MBIC11017 MASLPDFRPKQLSLGPLEFEILNLIWDLGTATVKQVHEQILTNPDRELAYTSVTTVLNRL
Anabaena7120 MAPLPDYRPKQMSVGPLEAEILNIVWEVGSATVKDVHDRILADPNRELAYTSVTTVLRRL
Bacillus_licheniformis_BlaI MKKIP-------QISDAELEVMKVIWKHSSINTNEVIKEL--SKTSTWSPKTIQTMLLRL
Staphylococcus_MecI ------MDNKTYEISSAEWEVMNIIWMKKYASANNIIEEI--QMQKDWSPKTIRTLITRL
Staphylococcus_BlaI ------MANKQVEISMAEWDVMNIIWDKKSVSANEIVVEI--QKYKEVSDKTIRTLITRL
 .:. * ::: ::* .. : : . : :: *:: **
Leptolyngbya_sp_PCC_7375 EKKGWIAKRR----VGRAY--HWEPLLSANNAKILESHERLHQFLAASNPDIVAAFADTL
Pseudoanabaena_sp_PCC_6802 VKKGWLTSHK----RGKIL--FWQAAISRPEAQALEAHERLNRFLEVGNADIVAAFADEL
Synechocystis TKKGWLVCHR----QGKAF--IWTARVSADQAKAVQSYEQLQQFLAISNPDVVAAFADSL
Synechococcus_sp_JA-2-3B'a(2-13) SQKGWVSCERRTDPQGRNLPMLWRPRLSRQEAASLRAFTHLQQFLAVGDPDTVAAFADSL
Acaryochloris_marina_MBIC11017 TKKGWLACDK----QNRSF--VWRPLVSRAQANSLWAYDQLQQFLAVGNPDTVAAFADHL
Anabaena7120 TDKGWLACDK----KGRAF--YWRPLLSKQQAQVIKAHDQLHSFLAVGNPDVVAAFADSL
Bacillus_licheniformis_BlaI IKKGALNHHK----EGRVF--VYTPNIDESDYIEVKSHSFLNRFYNGTLNSMVLNFLEN-
Staphylococcus_MecI YKKGFIDRKK----DNKIF--QYYSLVEESDIKYKTSKNFINKVYKGGFNSLVLNFVEK-
Staphylococcus_BlaI YKKEIIKRYK----SENIY--FYSSNIKEDDIKMKTAKTFLNKLYGGDMKSLVLNFAKN-
 .* : . . : . :. : : :: . . * * .
Leptolyngbya_sp_PCC_7375 DQDSLDQLEAITKRIQAARRARSNNQQGQEK
Pseud0anabaena_sp_PCC_6802 DLASVDRLEAIAQRLKAIRQEREER------
Synechocystis DTASIDQLTAIADRLRAARQQRQEEK-----
Synechococcus_sp_JA-2-3B'a(2-13) DQASVSQLQAIAERLRAARQARRDPR-----
Acaryochloris_marina_MBIC11017 DQASVEQLEEIADKIRAARKAREDQ------
Anabaena7120 DEAASEQIEAIAKRIQAARQAREEQ------
Bacillus_licheniformis_BlaI DQLSGEEINELYQILEEHKNRKKE-------
Staphylococcus_MecI EDLSQDEIEELRNILNKK-------------
Staphylococcus_BlaI EELNNKEIEELRDILNDISKK----------
 : . : : . :
Figure S7. Alignment of cyanobacterial PetR proteins with Staphylococcus and Bacillus BlaI/MecI proteins. Protein sequences were aligned using MUSCLE using the default parameters
Protein sequences were aligned using MUSCLE using the default parameters. The initial degradation site is highlighted in red.

### Slide 8
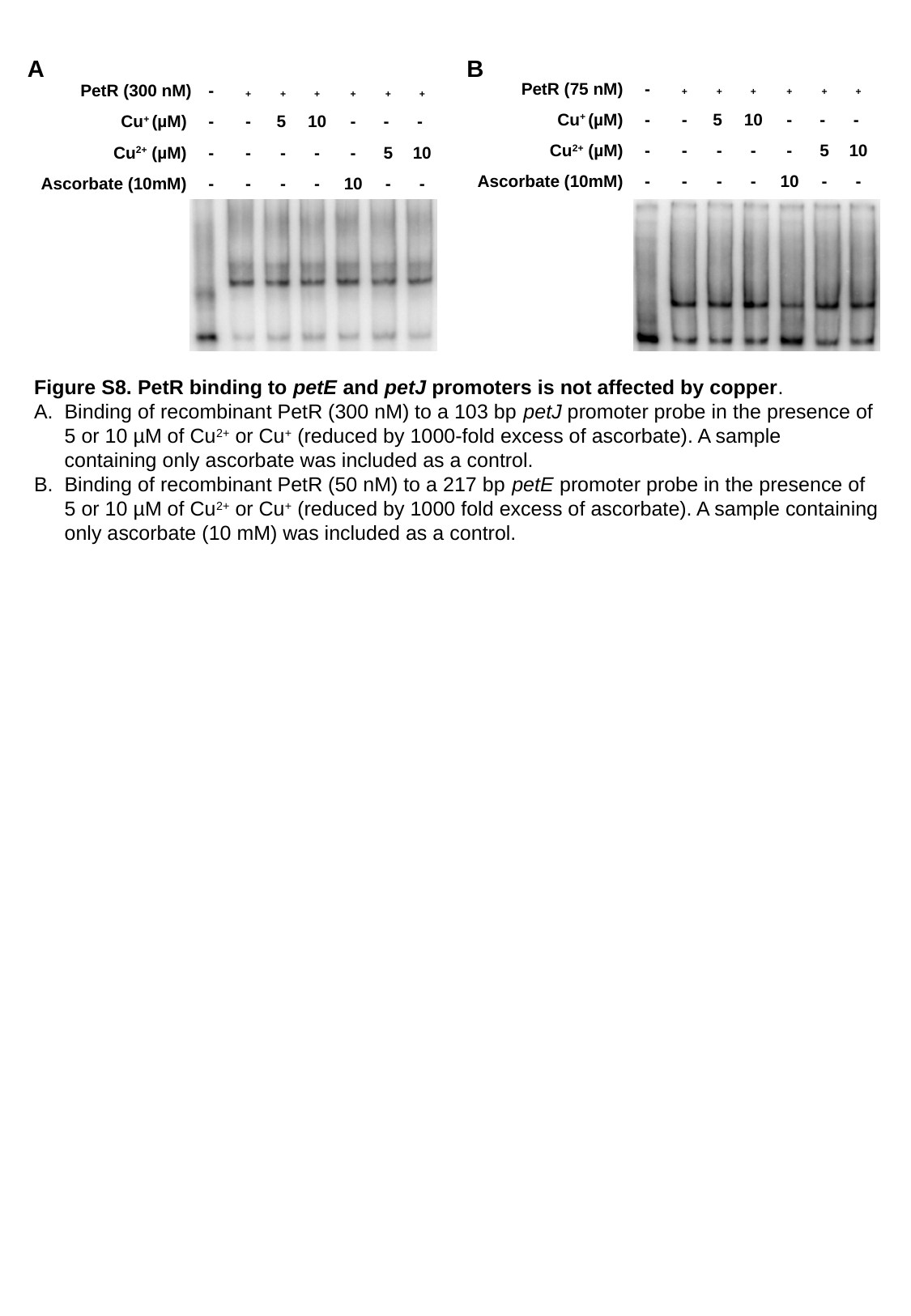

A
B
-
PetR (75 nM)
^+^
^+^
^+^
^+^
^+^
^+^
-
-
5
10
-
-
Cu^+^ (µM)
-
Cu^2+^ (µM)
-
-
-
-
-
5
10
-
-
-
-
10
-
-
Ascorbate (10mM)
PetR (300 nM)
-
^+^
^+^
^+^
^+^
^+^
^+^
-
-
5
10
-
-
Cu^+^ (µM)
-
Cu^2+^ (µM)
-
-
-
-
-
5
10
-
-
-
-
10
-
-
Ascorbate (10mM)
Figure S8. PetR binding to petE and petJ promoters is not affected by copper.
Binding of recombinant PetR (300 nM) to a 103 bp petJ promoter probe in the presence of 5 or 10 µM of Cu^2+^ or Cu^+^ (reduced by 1000-fold excess of ascorbate). A sample containing only ascorbate was included as a control.
Binding of recombinant PetR (50 nM) to a 217 bp petE promoter probe in the presence of 5 or 10 µM of Cu^2+^ or Cu^+^ (reduced by 1000 fold excess of ascorbate). A sample containing only ascorbate (10 mM) was included as a control.

### Slide 9
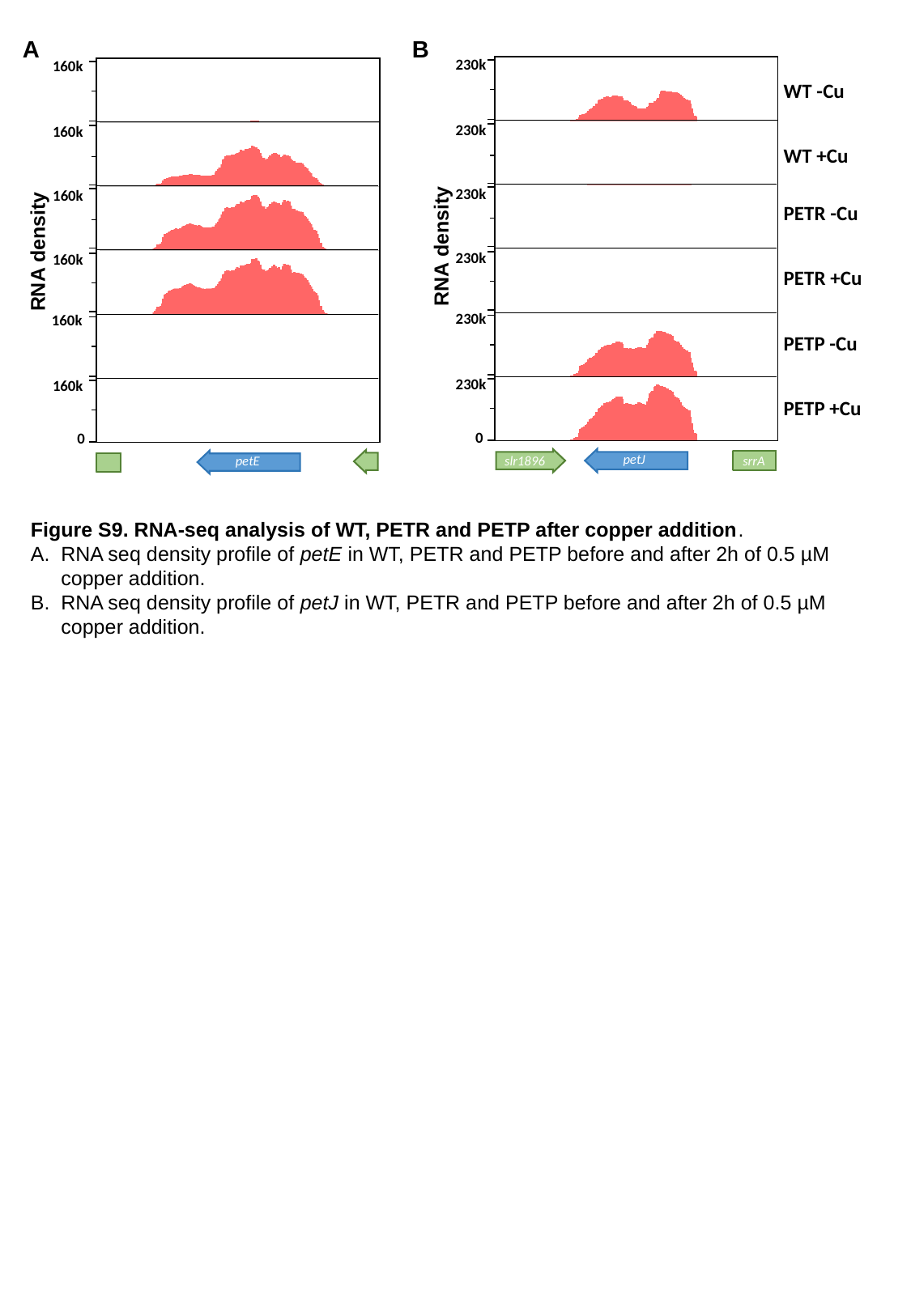

A
B
230k
230k
230k
230k
230k
230k
0
petJ
slr1896
srrA
WT -Cu
WT +Cu
PETR -Cu
PETR +Cu
PETP -Cu
PETP +Cu
RNA density
160k
160k
160k
RNA density
160k
160k
160k
0
petE
Figure S9. RNA-seq analysis of WT, PETR and PETP after copper addition.
RNA seq density profile of petE in WT, PETR and PETP before and after 2h of 0.5 µM copper addition.
RNA seq density profile of petJ in WT, PETR and PETP before and after 2h of 0.5 µM copper addition.

### Slide 10
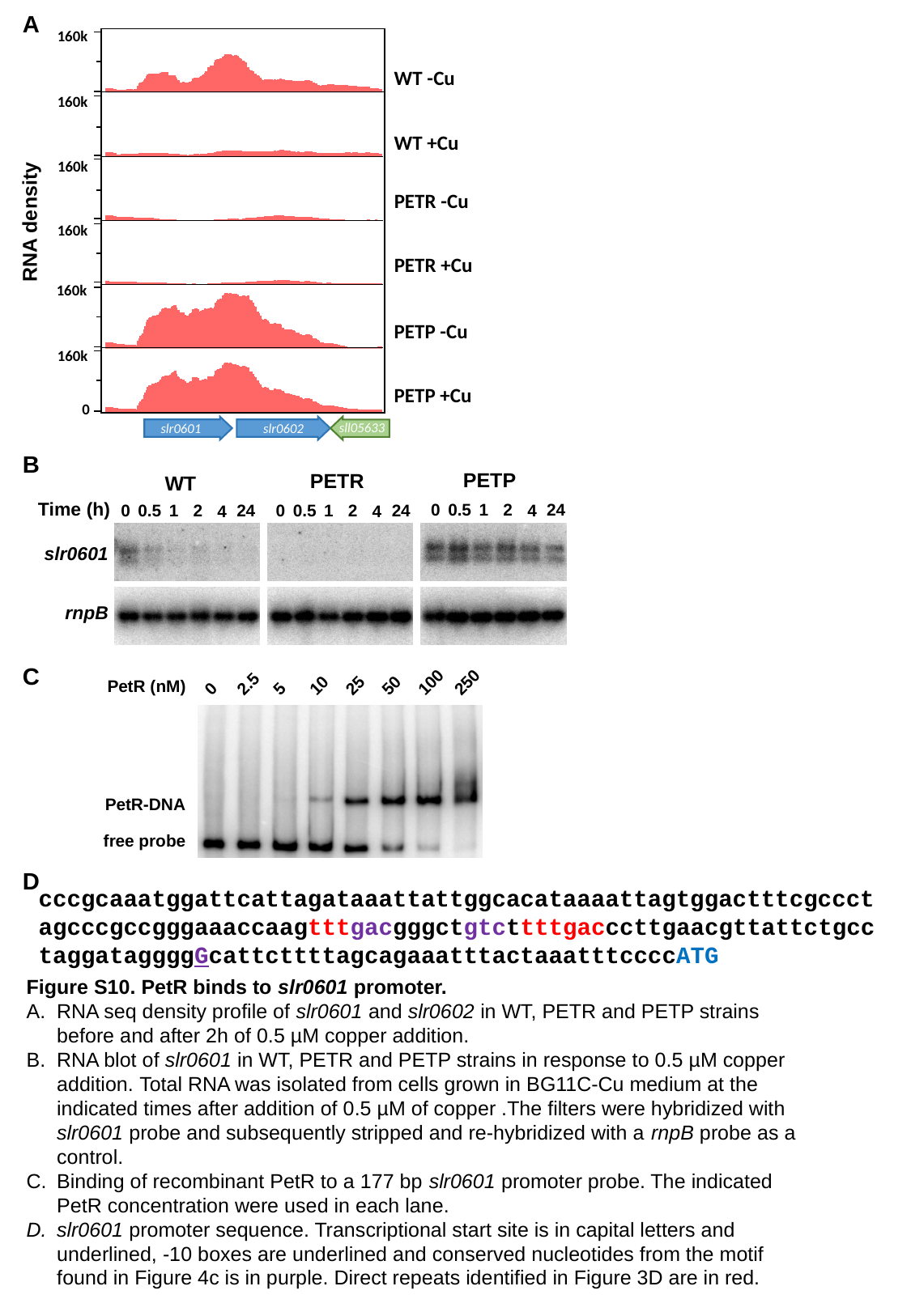

A
160k
160k
160k
RNA density
160k
160k
160k
0
sll05633
slr0601
slr0602
WT -Cu
WT +Cu
PETR -Cu
PETR +Cu
PETP -Cu
PETP +Cu
B
PETP
0
0.5
1
2
24
4
PETR
0
0.5
1
2
24
4
WT
0
0.5
1
2
24
4
Time (h)
slr0601
rnpB
C
0
2.5
5
10
25
50
100
250
PetR (nM)
PetR-DNA
free probe
D
cccgcaaatggattcattagataaattattggcacataaaattagtggactttcgccctagcccgccgggaaaccaagtttgacgggctgtcttttgacccttgaacgttattctgcctaggataggggGcattcttttagcagaaatttactaaatttccccATG
Figure S10. PetR binds to slr0601 promoter.
RNA seq density profile of slr0601 and slr0602 in WT, PETR and PETP strains before and after 2h of 0.5 µM copper addition.
RNA blot of slr0601 in WT, PETR and PETP strains in response to 0.5 µM copper addition. Total RNA was isolated from cells grown in BG11C-Cu medium at the indicated times after addition of 0.5 µM of copper .The filters were hybridized with slr0601 probe and subsequently stripped and re-hybridized with a rnpB probe as a control.
Binding of recombinant PetR to a 177 bp slr0601 promoter probe. The indicated PetR concentration were used in each lane.
slr0601 promoter sequence. Transcriptional start site is in capital letters and underlined, -10 boxes are underlined and conserved nucleotides from the motif found in Figure 4c is in purple. Direct repeats identified in Figure 3D are in red.

### Slide 11
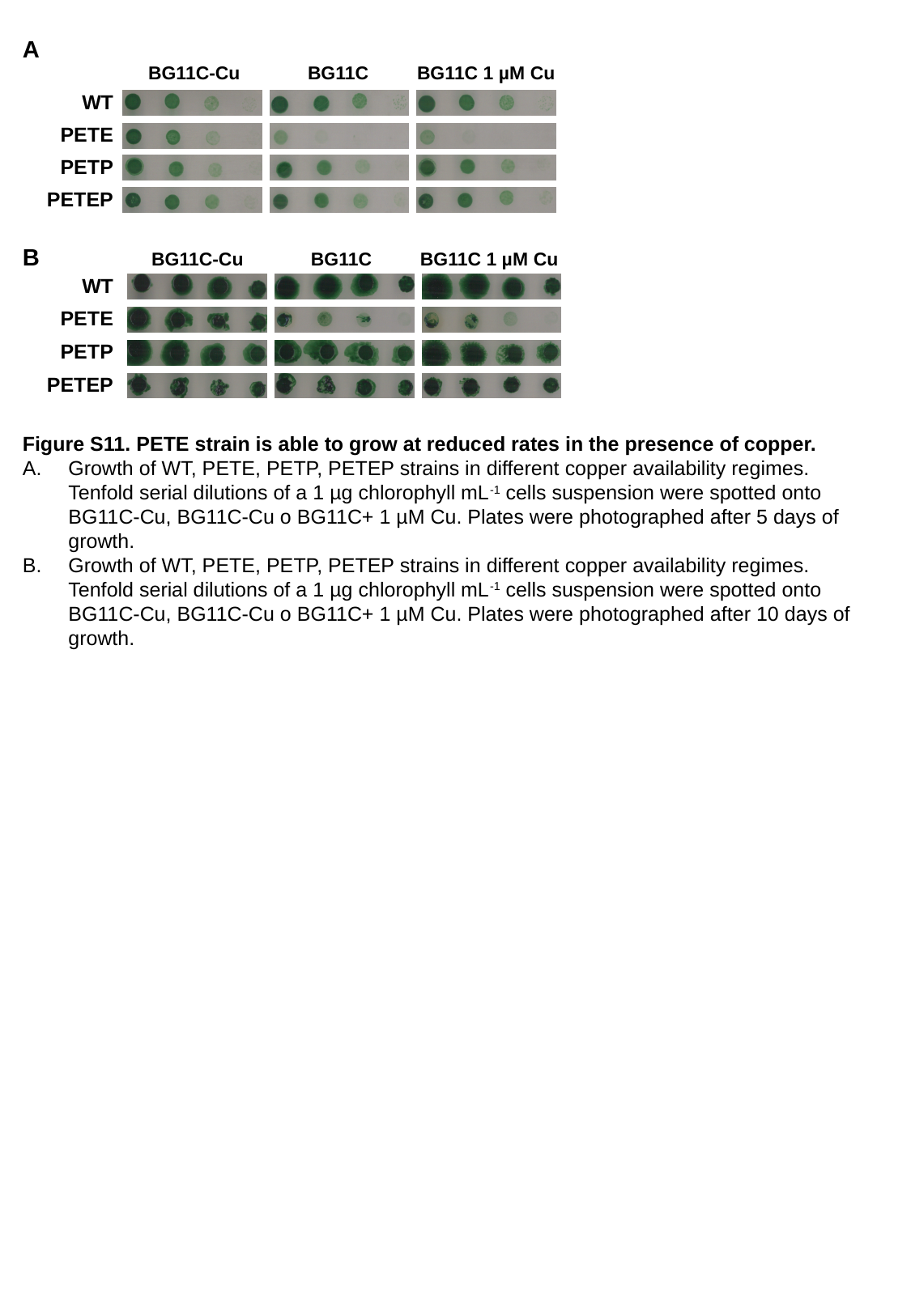

A
BG11C-Cu
BG11C
BG11C 1 µM Cu
WT
PETE
PETP
PETEP
B
BG11C-Cu
BG11C
BG11C 1 µM Cu
WT
PETE
PETP
PETEP
Figure S11. PETE strain is able to grow at reduced rates in the presence of copper.
Growth of WT, PETE, PETP, PETEP strains in different copper availability regimes. Tenfold serial dilutions of a 1 µg chlorophyll mL^-1^ cells suspension were spotted onto BG11C-Cu, BG11C-Cu o BG11C+ 1 µM Cu. Plates were photographed after 5 days of growth.
Growth of WT, PETE, PETP, PETEP strains in different copper availability regimes. Tenfold serial dilutions of a 1 µg chlorophyll mL^-1^ cells suspension were spotted onto BG11C-Cu, BG11C-Cu o BG11C+ 1 µM Cu. Plates were photographed after 10 days of growth.

### Slide 12
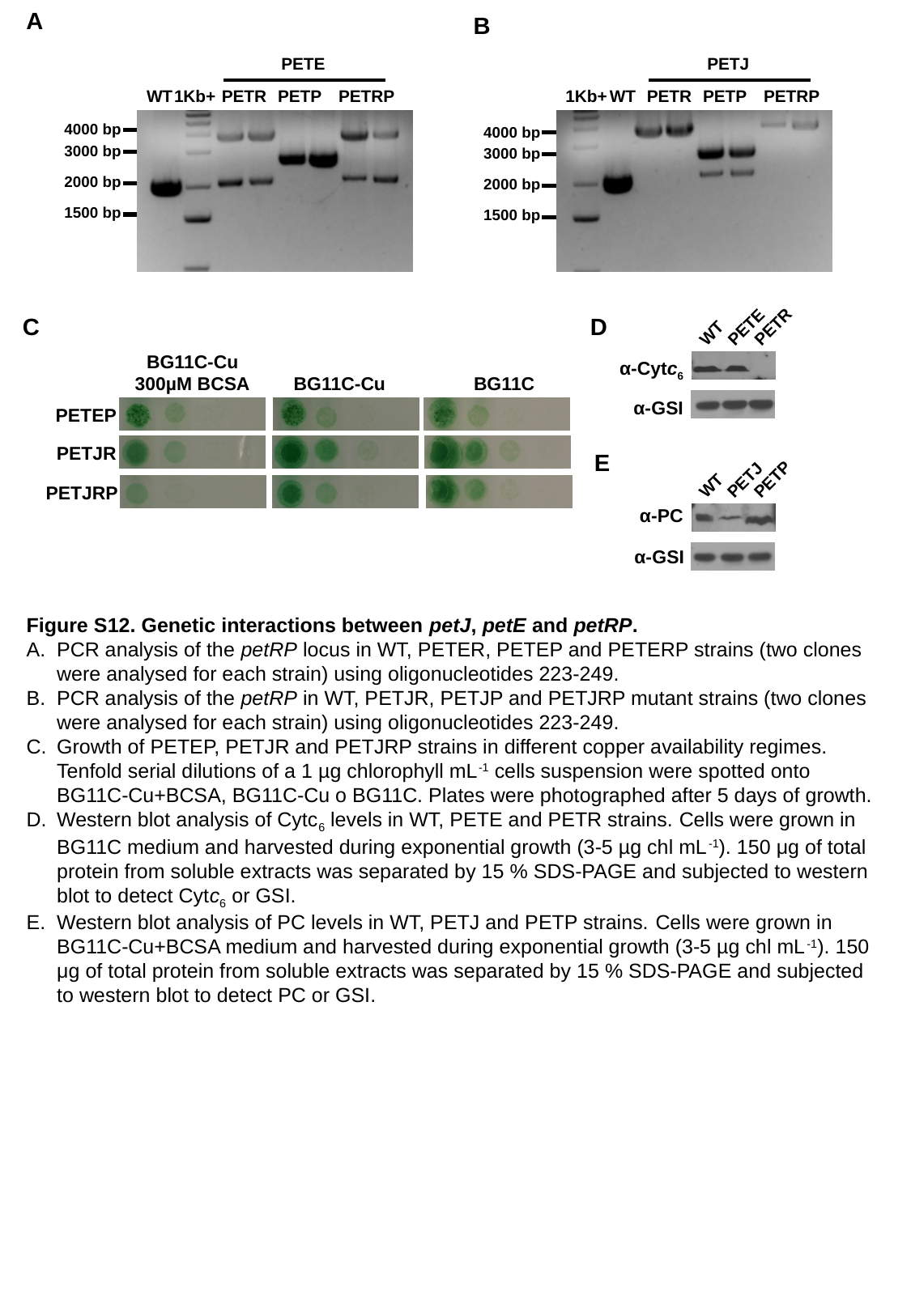

A
B
PETJ
1Kb+
WT
PETR
PETP
PETRP
4000 bp
3000 bp
2000 bp
1500 bp
PETE
WT
1Kb+
PETR
PETP
PETRP
4000 bp
3000 bp
2000 bp
1500 bp
C
D
WT
PETE
PETR
α-Cytc_6_
α-GSI
BG11C-Cu
300µM BCSA
BG11C-Cu
BG11C
PETEP
PETJR
PETJRP
E
WT
PETJ
PETP
α-PC
α-GSI
Figure S12. Genetic interactions between petJ, petE and petRP.
PCR analysis of the petRP locus in WT, PETER, PETEP and PETERP strains (two clones were analysed for each strain) using oligonucleotides 223-249.
PCR analysis of the petRP in WT, PETJR, PETJP and PETJRP mutant strains (two clones were analysed for each strain) using oligonucleotides 223-249.
Growth of PETEP, PETJR and PETJRP strains in different copper availability regimes. Tenfold serial dilutions of a 1 µg chlorophyll mL^-1^ cells suspension were spotted onto BG11C-Cu+BCSA, BG11C-Cu o BG11C. Plates were photographed after 5 days of growth.
Western blot analysis of Cytc_6_ levels in WT, PETE and PETR strains. Cells were grown in BG11C medium and harvested during exponential growth (3-5 µg chl mL^-1^). 150 μg of total protein from soluble extracts was separated by 15 % SDS-PAGE and subjected to western blot to detect Cytc_6_ or GSI.
Western blot analysis of PC levels in WT, PETJ and PETP strains. Cells were grown in BG11C-Cu+BCSA medium and harvested during exponential growth (3-5 µg chl mL^-1^). 150 μg of total protein from soluble extracts was separated by 15 % SDS-PAGE and subjected to western blot to detect PC or GSI.

### Slide 13
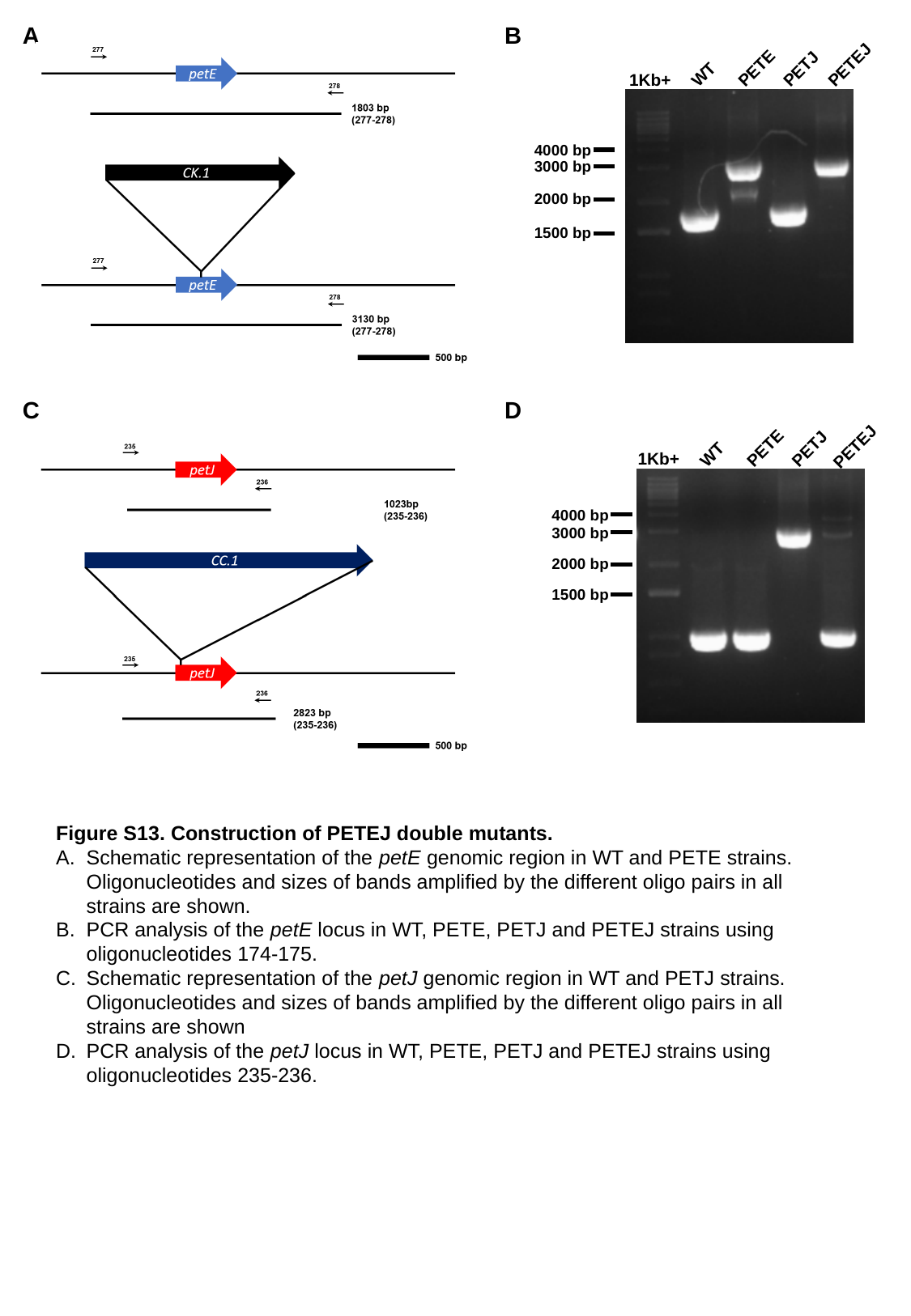

A
B
PETEJ
WT
PETE
PETJ
1Kb+
4000 bp
3000 bp
2000 bp
1500 bp
C
D
PETEJ
WT
PETE
PETJ
1Kb+
4000 bp
3000 bp
2000 bp
1500 bp
Figure S13. Construction of PETEJ double mutants.
Schematic representation of the petE genomic region in WT and PETE strains. Oligonucleotides and sizes of bands amplified by the different oligo pairs in all strains are shown.
PCR analysis of the petE locus in WT, PETE, PETJ and PETEJ strains using oligonucleotides 174-175.
Schematic representation of the petJ genomic region in WT and PETJ strains. Oligonucleotides and sizes of bands amplified by the different oligo pairs in all strains are shown
PCR analysis of the petJ locus in WT, PETE, PETJ and PETEJ strains using oligonucleotides 235-236.

### Slide 14
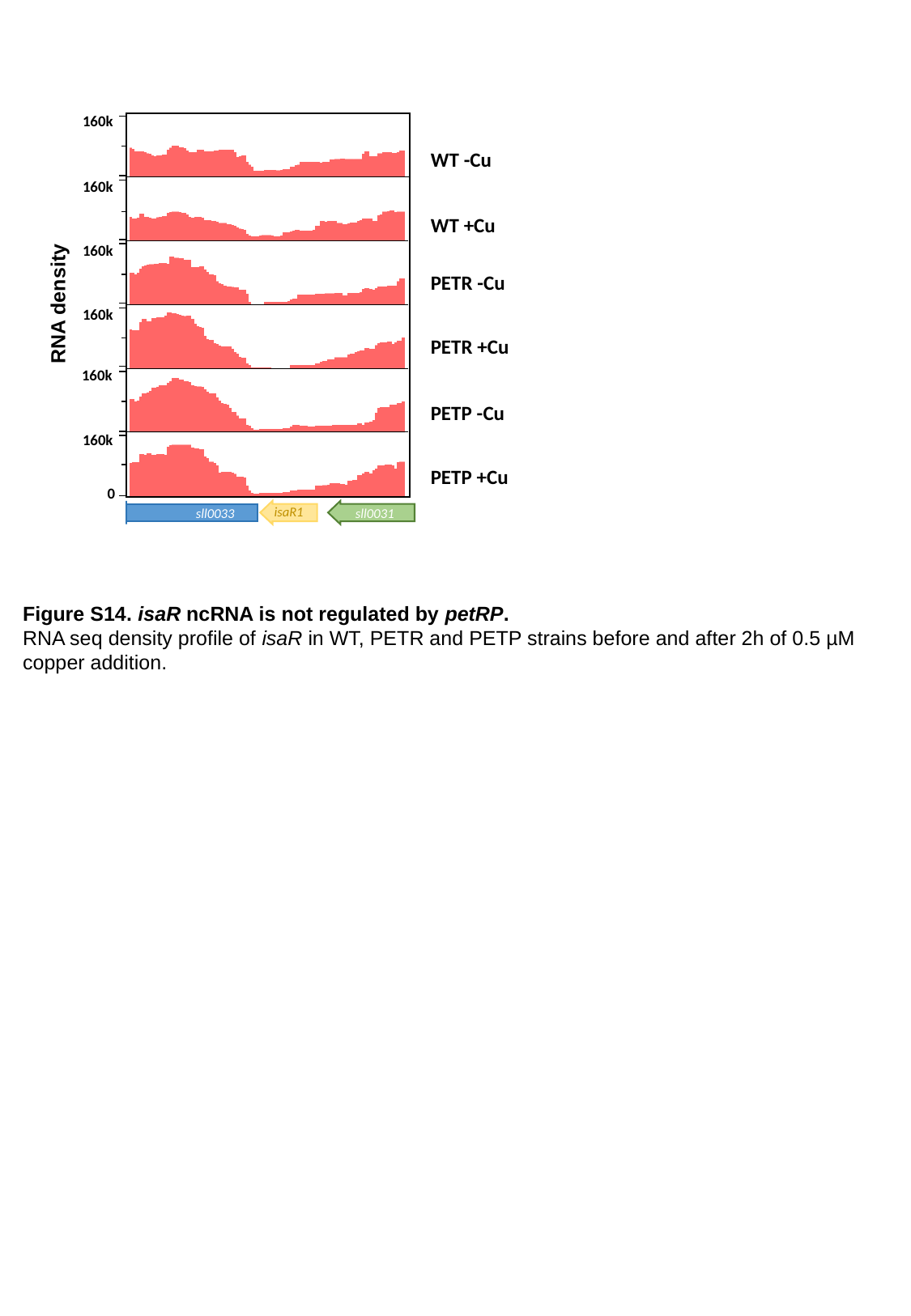

160k
WT -Cu
WT +Cu
PETR -Cu
PETR +Cu
PETP -Cu
PETP +Cu
160k
160k
RNA density
160k
160k
160k
0
isaR1
sll0033
sll0031
Figure S14. isaR ncRNA is not regulated by petRP.
RNA seq density profile of isaR in WT, PETR and PETP strains before and after 2h of 0.5 µM copper addition.

### Slide 15
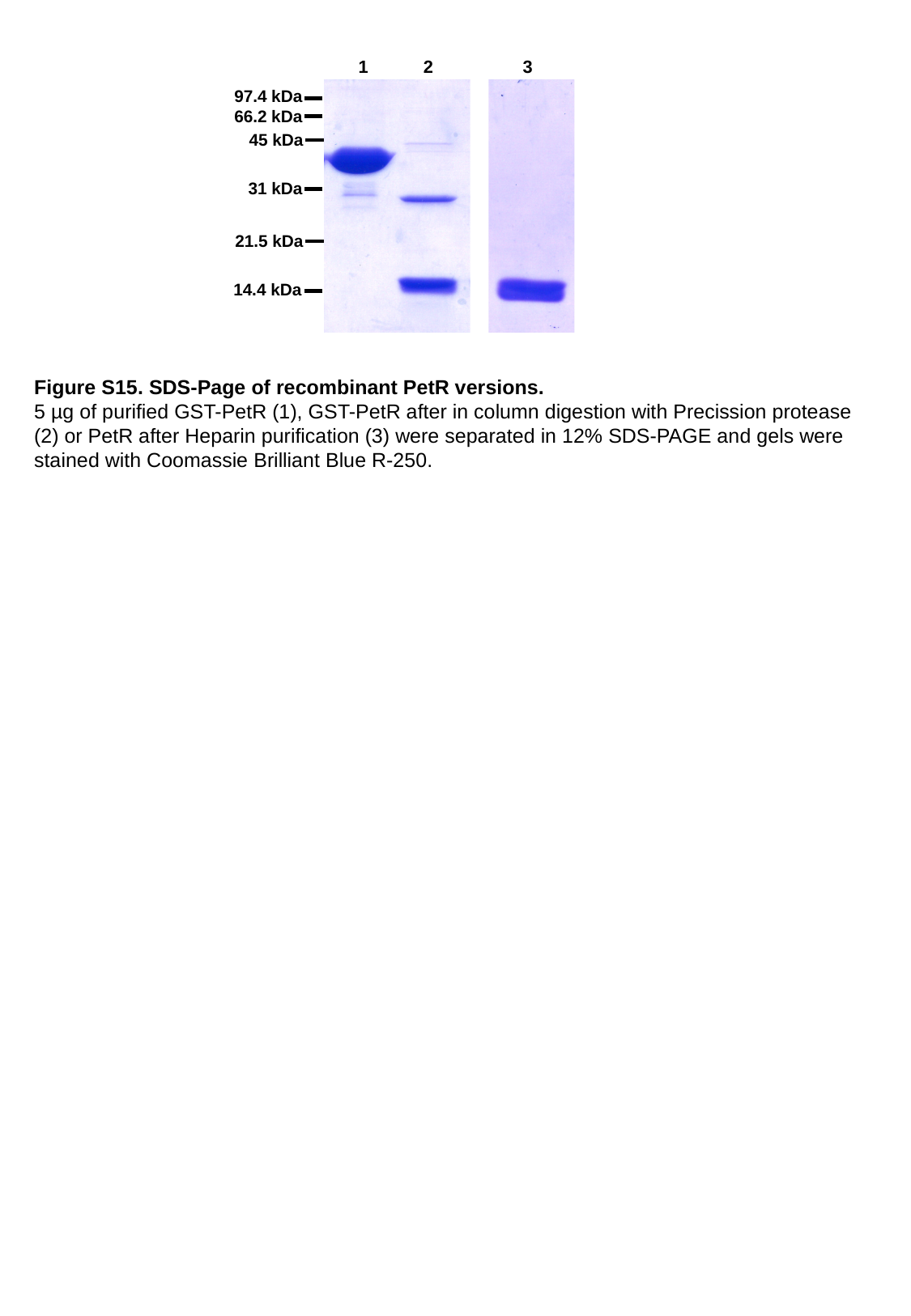

1
2
3
97.4 kDa
 66.2 kDa
45 kDa
31 kDa
21.5 kDa
14.4 kDa
Figure S15. SDS-Page of recombinant PetR versions.
5 µg of purified GST-PetR (1), GST-PetR after in column digestion with Precission protease (2) or PetR after Heparin purification (3) were separated in 12% SDS-PAGE and gels were stained with Coomassie Brilliant Blue R-250.
